# Individualized parcellation reveals functional boundaries in human prefrontal cortex

**DOI:** 10.64898/2026.07.04.736504

**Authors:** Jinkang Derrick Xiang, Da Zhi, Bassel Arafat, Caroline Nettekoven, Jörn Diedrichsen, Marieke Mur

## Abstract

Human prefrontal cortex (PFC) supports diverse higher-order cognitive functions during task execution. Increasing evidence suggests that these functions are organized along large-scale gradients, such as a rostro-caudal axis supporting progressively abstract-to-concrete information processing. At the same time, regional specialization within PFC, including focal patches selective for specific stimulus categories, suggests the presence of discrete functional boundaries. Whether PFC organization is best characterized by continuous gradients or discrete subdivisions remains unresolved. Here we show that task-evoked functional organization in PFC exhibits substantial inter-individual variability, limiting the usefulness of conventional group atlases in testing for functional boundaries. To address this challenge, we estimated individualized functional parcellations by combining a group atlas with task-evoked fMRI data spanning diverse cognitive tasks. The group atlas revealed large-scale functional gradients across PFC, whereas individualized parcellations additionally uncovered sharp functional boundaries obscured by group averaging. Moreover, functional organization in PFC was substantially more fine-grained than in other association cortices, consistent with its integrative role in cognitive control. Together, these findings suggest that PFC organization reflects an individualized mosaic of fine-grained functional subdivisions embedded within broader large-scale gradients, with important implications for defining core constructs such as the multiple-demand system.

## 1. Introduction

Primate prefrontal cortex (PFC) is central to cognition, supporting diverse functions such as working memory, decision-making, prospective memory, and motor planning (Duncan, 2001, Miller and Cohen, 2001, Duncan, 2010). Yet the functional organization of PFC remains debated. A recent large-scale meta-analysis of task-evoked activity patterns showed the existence of continuous functional gradients across PFC (Abdallah et al., 2022), consistent with earlier accounts of rostro–caudal transitions from abstract to concrete control and ventro–dorsal differentiation related to perceptual–motor transformations and semantic processing (Badre, 2008, O’Reilly, 2010). Together, these findings suggest a smoothly varying functional architecture.

However, empirical work also reveals spatially delineated clusters of functional specialization within PFC, suggesting the presence of functional boundaries rather than purely continuous gradients. For example, in macaque PFC, cortical patches have been identified that selectively respond to faces (Tsao et al., 2008) as well as auditory and audiovisual information (Romanski and Goldman-Rakic, 2002, Sugihara et al., 2006). These findings raise the question of whether such functional boundaries are present throughout human PFC or whether its overall organization is better characterized as graded (Abdallah et al., 2022, Badre, 2008, O’Reilly, 2010).

A central challenge in identifying functional boundaries in PFC is the substantial inter-individual variability in its functional organization. Even after accounting for anatomical differences, functional brain organization varies markedly across individuals, particularly in the association cortex (Mueller et al., 2013, Gordon et al., 2017b, Gratton et al., 2018, Dworetsky et al., 2024). In humans, functional brain parcellations typically divide cortex into spatially contiguous regions mainly based on resting-state functional connectivity patterns (Thomas Yeo et al., 2011, Schaefer et al., 2018, Ji et al., 2019, Glasser et al., 2016). These atlases, however, are derived from group-averaged data and therefore assume a common spatial layout across individuals. This raises the possibility that fine-grained boundaries present in individual brains may be obscured at the group level. Critically, this issue has not been systematically evaluated using a broad set of task-evoked responses, which have been shown to provide a better prediction of functional organization than resting-state correlations (Nettekoven et al., 2026).

We therefore investigated the functional organization of PFC using six richly sampled multi-task fMRI datasets in which a smaller number of participants performed a broad array of cognitive tasks (King et al., 2019, Pinho et al., 2018, Nakai and Nishimoto, 2020, Barch et al., 2013, Assem et al., 2024, Arafat et al., 2026). We first show that task-evoked response profiles in PFC vary substantially across individuals, motivating individualized mapping. For each participant, we therefore derive an individualized functional parcellation by combining a group atlas with subject-specific task-evoked activity (Zhi et al., 2025, Nettekoven et al., 2024). We then test whether the group and individualized atlases capture sharp transitions in response preferences using a distance-controlled boundary coefficient (DCBC) (Zhi et al., 2022, King et al., 2019). We show that task-evoked response similarity in PFC declines with spatial distance, consistent with graded organization. Parcel borders in group atlases do not capture additional structure beyond this smooth spatial variation. In contrast, individualized parcellations reveal significant functional boundaries. Furthermore, we demonstrate that the spatial layout of task-evoked activity in PFC is organized at a finer spatial scale than parietal cortex. Together, these findings indicate that human PFC exhibits a fine-grained and individualized functional architecture in which discrete boundaries coexist with continuous gradients. These organizational principles motivate a re-evaluation of core constructs of PFC function, which often are defined using group-averaged activity maps. As an example, we re-evaluate the idea that different executive functions all activate a common core multiple-demand system (Duncan, 2010, Assem et al., 2020).

## 2. Results

### 2.1. Task-evoked activity in PFC exhibits substantial inter-individual variability

To study the functional boundaries in individual participants, we leveraged six large-scale fMRI datasets (see Supplementary Table 1), in which participants performed a series of higher-order cognitive tasks spanning multiple domains, including cognitive, social, motor, affective and language processing (King et al., 2019, Pinho et al., 2018, Assem et al., 2024, Barch et al., 2013, Nakai and Nishimoto, 2020, Arafat et al., 2026), sampled across a large number of task conditions. In each dataset, the task-evoked activities are the regression weights associated with each task condition, normalized by the noise standard deviation for each voxel (see methods). Thus for each vertex and individual, we obtain a functional profile, the vector of task-evoked activity across tasks.

As a first step, we aimed to quantify how much inter-individual variability in the functional organization there is. This is straightforward in a task-based imaging approach. For each dataset, we decomposed the variance of the functional profile across subjects into three components: a component that is consistent across subjects (group, *g*), a component that is unique to each subject and replicable across runs (subject, *s*), and a component that fluctuates across runs (noise, *ε*) (Seghier and Price, 2018, Ejaz et al., 2015, King et al., 2019). The components are orthogonal to each other and capture the total variability in task-evoked neural activity. This variance decomposition can be done vertex-wise or for the multi-vertex response pattern. The variance explained by the group signal (*V_g_*) and the subject signal (*V_s_*) constitutes the explainable variance (*V_g_* + *V_s_*). The amount of explainable variance (functional signal-to-noise ratio) varies across brain regions. To quantify inter-individual variability, we computed the proportion of subject-specific variance out of the explainable variance, i.e., *V_s_/*(*V_g_* + *V_s_*).

We first performed variance decomposition at each vertex to generate whole-cortex maps of inter-individual variability (Fig. 1a). We found the lowest inter-subject variability in lower-level visual areas and primary auditory cortex. In contrast, association areas in the temporal, parietal, and especially prefrontal cortices exhibited very high inter-individual variability, with more than 60% of reliable variance being idiosyncratic.

**Figure 1:**
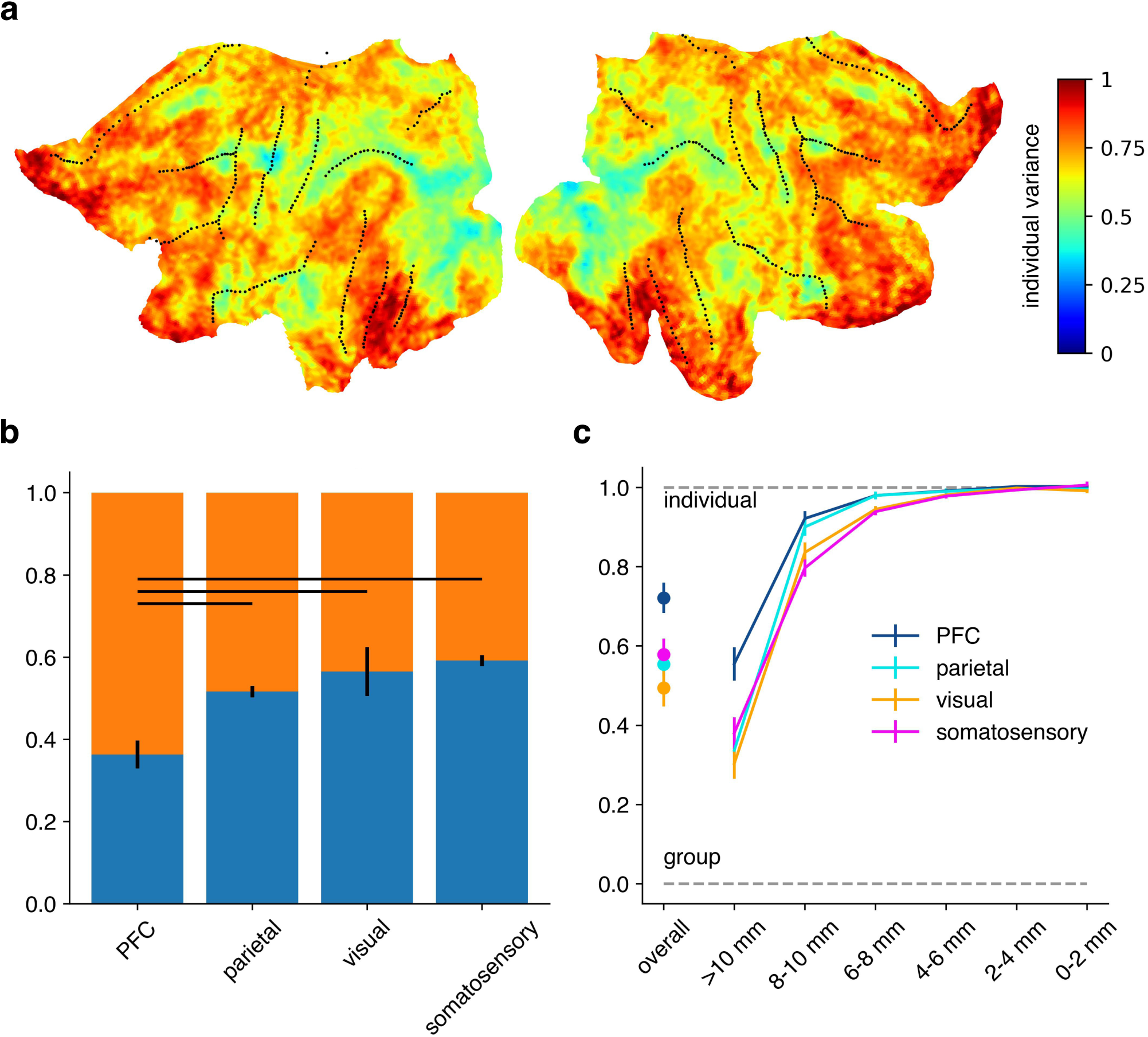
Decomposition of task-evoked variance into group and individual components. (**a**) Variance decomposition for the left and right hemispheres across 6 datasets. Values indicate the proportion of subject-specific variance out of explainable variance for each vertex. Zero means that functional profiles are the same across subjects, one means that functional profiles are fully idiosyncratic (uncorrelated across subjects). **(b)** Variance decomposition averaged in 4 regions of interest for both hemispheres. The height of the blue bars show the average proportion of group variance out of explainable variance, the orange bars show the average proportion of subject-specific variance. Error bars show standard error across subjects and datasets (*n* = 146). Horizontal lines indicate significant differences (*p <* 0.05, uncorrected, mixed linear model). **(c)** Variance decomposition across spatial frequencies averaged in 4 regions of interest across datasets. Values indicate the proportion of subject-specific variance out of explainable variance. Error bars show standard error across datasets (*n* = 6).

For regional comparisons and statistical inference, we next applied variance decomposition to the multivariate response patterns in regions of interest (ROIs) (Fig. 1b,c). We fit a linear mixed-effects model with ROIs as fixed effects and datasets as random effects. Using PFC as the reference category, we found that PFC exhibited significantly greater individual variance than parietal, somatosensory, and visual cortex across all six datasets. Relative to

PFC, the proportion of individual variance was lower in parietal cortex (*β* = −0.153, *p* = 0.002), somatosensory cortex (*β* = −0.202, *p <* 0.001), and visual cortex (*β* = −0.229, *p <* 0.001). The intercept estimate for PFC was 0.672 (*p <* 0.001), indicating that approximately 67% of explainable variance in PFC was subject-specific.

Furthermore, the estimated variance associated with the random dataset effect was small (*σ*^2^ = 0.006), suggesting that the regional differences generalized consistently across datasets. The subset of Glasser parcels used to define each ROI is listed in Supplementary Table 2.

We further hypothesized that the inter-individual variability mostly affects the detailed layout of the functional organization, while the broad functional gradients should remain quite reliable across subjects. To test this idea, we decomposed each individual functional map into a series of maps capturing the smoothed overall gradients up to the finest spatial details at the vertex-to-vertex level (see Methods 4.2). Within each frequency band, we then performed the decomposition into group and individual variance (King et al., 2019). Consistent with our hypothesis, we found that the group contribution (correlation across subjects) was largest for maps smoothed with 10 mm kernels. For finer spatial details, there was less group topology (i.e., more idiosyncrasies). At a scale finer than 6 mm/cycle, we found hardly any group topology, with subject-specific signal dominating these finer spatial scales (Fig. 1c). Overall, we found that the functional organization in PFC was idiosyncratic. While there is clearly a common organization in the coarser spatial gradients, the finer details of the functional organization were highly idiosyncratic.

### 2.2. Group atlas provides no evidence for functional boundaries in PFC

Given the high degrees of idiosyncrasies of the fine-grained functional organization, group parcellations may have limited power to detect reliable functional boundaries in PFC.

To test this idea, we tested whether the regions defined by the Glasser atlas (Glasser et al., 2016) would align to individual functional boundaries in PFC. To do so, we computed a distance-controlled boundary coefficient (DCBC) (Zhi et al., 2022) on the MDTB dataset (King et al., 2019), using the functional profile of each voxel. DCBC is defined as the difference of within-parcel and between-parcel correlations of the functional profiles among vertices. The correlations are estimated with cross-validation to eliminate the influence of measurement noise. Finally, to account for the intrinsic spatial smoothness of the data, DCBC only compares pairs of vertices with the same distance on the cortical surface (Methods 4.4). Thus, if PFC had only smooth functional gradients but no functional boundaries, then DCBC should be close to 0.

In PFC, the correlation of functional profiles decreased with increasing spatial distance (Fig. 2c), consistent with the presence of overall functional gradients. However, the boundaries defined by the Glasser group atlas (Glasser et al., 2016) did not align with places where the functional profiles changed more rapidly (i.e. with functional boundaries), which is evident by no significant differences of the averaged correlations of within– and between-parcel vertex pairs in each spatial bin (Fig. 2c). For statistical comparison, we derived a single evaluation measure, where DCBC is weighted averaged across spatial bins (Zhi et al., 2022) up to 35 mm, and we found the average DCBC value for the Glasser group atlas was not significantly greater than 0 (one-sided t-test, *t*(23) = 0.089*, p* = 0.465) as in Fig. 2c,e, suggesting the group atlas does not reveal functional boundaries of individual PFC.

**Figure 2:**
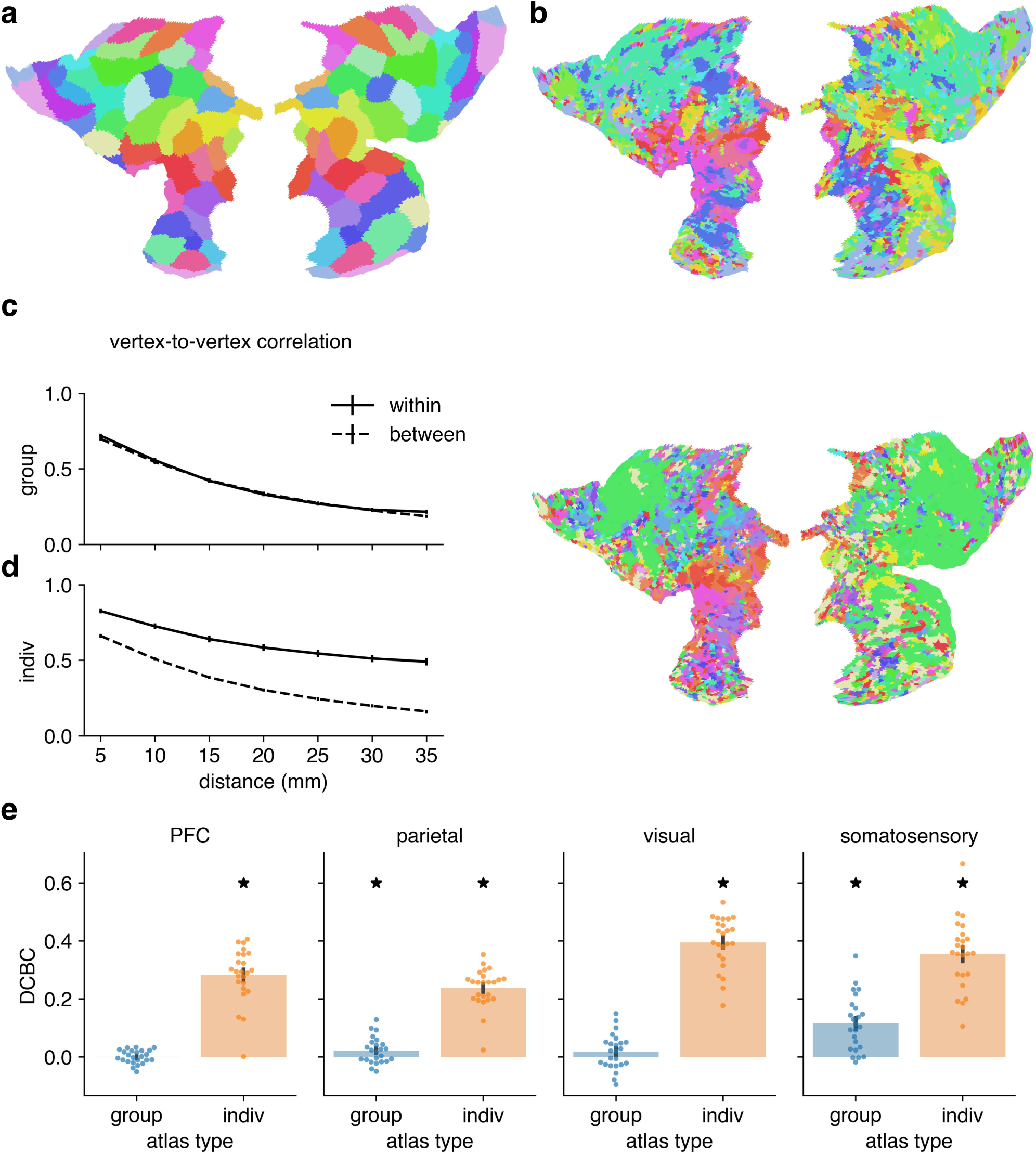
DCBC Glasser. (**a**) Glasser group atlas for PFC (Glasser et al., 2016). **(b)** Individualized Glasser atlas for two example subjects. **(c)** Average cross-validated correlation as a function of spatial distance for functional parcel boundaries for the Glasser group atlas. The DCBC is defined as the difference in correlation (within – between) within each distance bin. The error bars show standard error across participants (*n* = 24). **(c)** As in **(c)**, but for the individualized Glasser atlas. **(e)** Average DCBC for the Glasser group (blue) and individualized (orange) atlases across subjects (*n* = 24). Error bars show standard error across subjects. Asterisks indicate that DCBC is significantly higher than 0 (one-sided t-test, *p <* 0.05, uncorrected). For each ROI, DCBC on the individualized atlas is significantly higher than the group (two-sided t-test, *p <* 0.05, uncorrected, significance not shown).

These results were replicated with the Schaefer atlas (see Supplementary Fig. 1a,c).

### 2.3. Individualized parcellation reveals functional boundaries in PFC

Given the substantial inter-individual variability of neural activities in PFC, we reasoned that real functional boundaries, if they exist, would likely only become visible if we can derive cortical parcellations for single subjects (Eickhoff et al., 2018). However, it typically requires an extensive amount of individual functional data to derive a reliable functional characterization (Marek et al., 2018), which is prohibitive. We leveraged a hierarchical Bayesian brain parcellation framework (Zhi et al., 2025) to combine a group atlas with limited individual functional data to derive an individualized parcellation for each subject in the MDTB dataset.

More specifically, we derived an individualized parcellation *U^s^* for each subject by combining their functional data with a group-level prior using the hierarchical Bayesian framework (Zhi et al., 2025). This approach allows parcel boundaries to shift according to each subject’s task-evoked activity (captured by response profiles), while remaining anchored to the overall organizational structure captured at the group level (Zhi et al., 2025, Nettekoven et al., 2024). As a result, the individualized parcellations reveal subject-specific functional organization that is not apparent in the group atlas. Each cortical location was assigned to the parcel with the highest posterior probability, yielding a subject-specific parcellation (Fig. 2b). This subject-specific refinement enabled us to test for functional boundaries within individuals.

We trained the individualized parcellation using one set of task conditions of the MDTB dataset and evaluated it in a cross-validated way using another set of task conditions. The two task sets sample tasks of similar domains but differ in tasks and conditions. The differences between averaged correlation of within– and between-parcel vertex pairs of the individualized atlases is much larger than those of the group atlases in each spatial bin (Fig. 2c,d), suggesting that the individualized parcellations can detect real task-invariant functional boundaries within each individual.

We then computed the DCBC value for each ROI based on group and individual parcellation. DCBC is significantly higher on the individualized Glasser atlas than the group atlas for all the ROIs (two-sided t-tests, uncorrected; PFC: *t*(46) = −14.586*, p <* 0.001; parietal: *t*(46) = −12.775*, p <* 0.001; visual: *t*(46) = −17.340*, p <* 0.001; somatosensory: *t*(46) = −7.511*, p <* 0.001). In addition, DCBC is significantly above 0 on the individualized Glasser atlas for all the ROIs (one-sided t-tests, uncorrected; PFC: *t*(23) = 15.057*, p <* 0.001; parietal: *t*(23) = 16.850*, p <* 0.001; visual: *t*(23) = 21.972*, p <* 0.001; somatosensory: *t*(23) = 14.247*, p <* 0.001). Results shown in Fig. 2e.

Overall, the Glasser group atlas only suggests functional gradients in PFC. The individualized atlas, taking into account the idiosyncrasies of task-evoked neural activity, highlights the existence of functional boundaries.

Results are qualitatively similar for the Schaefer atlas (Schaefer et al., 2018), as in Supplementary Fig. 1.

### 2.4. PFC shows a finer spatial scale of functional organization than parietal cortex

Our results so far suggest that the PFC is organized as a mosaic of functional regions that are spatially arranged with a high level of inter-individual variability. This raises the question whether this is also true for other associative cortical areas. As a comparison we chose the posterior parietal cortex. The prefrontal cortex and the parietal cortex carry out similar functions in higher-order cognition and have extensive connections with each other (Goldman-Rakic, 1984, Xu et al., 2022). Are putative functional regions organized at similar or different spatial scales?

To quantitatively test this, we computed the spatial correlation between the functional profile for each vertex with its surrounding neighbors. To remove the influence of spatially auto-correlated measurement noise, this correlation was determined in a cross-validated fashion. We found that the similarity of functional profiles dropped off in both regions (Fig. 3a). For the same distances on the cortical surface, however, the functional profile changes more rapidly in the PFC than in the parietal cortex.

**Figure 3:**
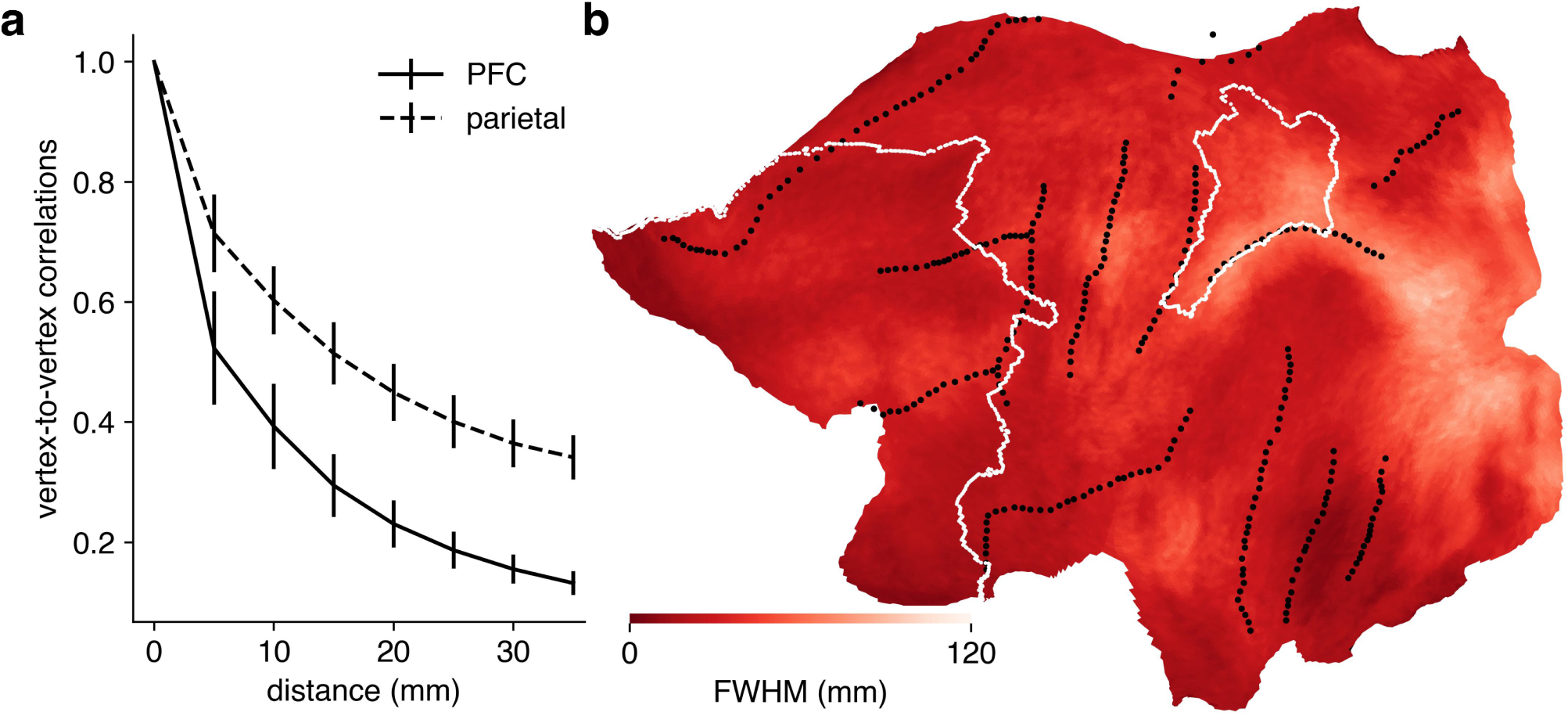
Fine-grained functional organization in PFC. (**a**) Cross-validated spatial autocorrelation function of the response profiles for PFC and parietal regions. Error bars show standard error across datasets (*n* = 6). **(b)** Full-width-at-half-maximum (FWHM) of the spatial autocorrelation function for each vertex, averaged across subjects and datasets (*n* = 146).

For visualization purposes, we estimated the full-width-at-half-maximum (FWHM) of the spatial autocorrelation function for each vertex and plotted on the surface. Overall vertices in PFC have lower FWHMs compared to parietal and visual cortices, suggesting a fine-grained functional organization (Fig. 3b).

### 2.5. Group averaging overestimates functional overlap in the multiple-demand system

Our results indicate that the functional organization in the prefrontal cortex is highly idiosyncratic, fragmented and structured by functional boundaries. These observations have implications for how the functional organization should be studied.

One of the core constructs in theories of PFC function is the multiple-demand (MD) system (Duncan, 2010). The MD system refers to a set of frontal and parietal regions that are recruited across diverse cognitive demands and exhibit domain-general responses that scale with the difficulty of the task (Duncan, 2001, 2010). Evidence for such domain-general responses has largely been derived from group-averaged data (Duncan and Owen, 2000, Duncan, 2001, 2010), which may overestimate the degree of overlap across tasks (Fedorenko et al., 2013).

Here, we tested whether overlap between executive tasks in prefrontal activity patterns was also present at the level of individual subjects. To do so, we re-analyzed the data from Assem et al. (2024), comparing correlations between activity patterns for the N-back, task-switching, and stop-signal tasks estimated from group-averaged data with those estimated within individual subjects. Specifically, for each task we computed the activity contrast between the hard and easy conditions (Assem et al., 2024), and then quantified the similarity of these contrast patterns across tasks within core frontal MD regions.

Because activity patterns estimated in individual subjects are substantially noisier than group-averaged maps, directly comparing raw correlations would systematically underestimate correspondence at the individual level. We therefore used a noise-corrected correlation measure that estimates the correlation between the underlying true activity patterns after accounting for measurement noise (Diedrichsen et al., 2026). In parallel, we estimated the functional signal-to-noise ratio (fSNR), defined as the ratio between the variance of the true response patterns and measurement noise (Diedrichsen et al., 2026), to ensure that differences in correlation were not driven by low signal quality (Fig. 4).

**Figure 4:**
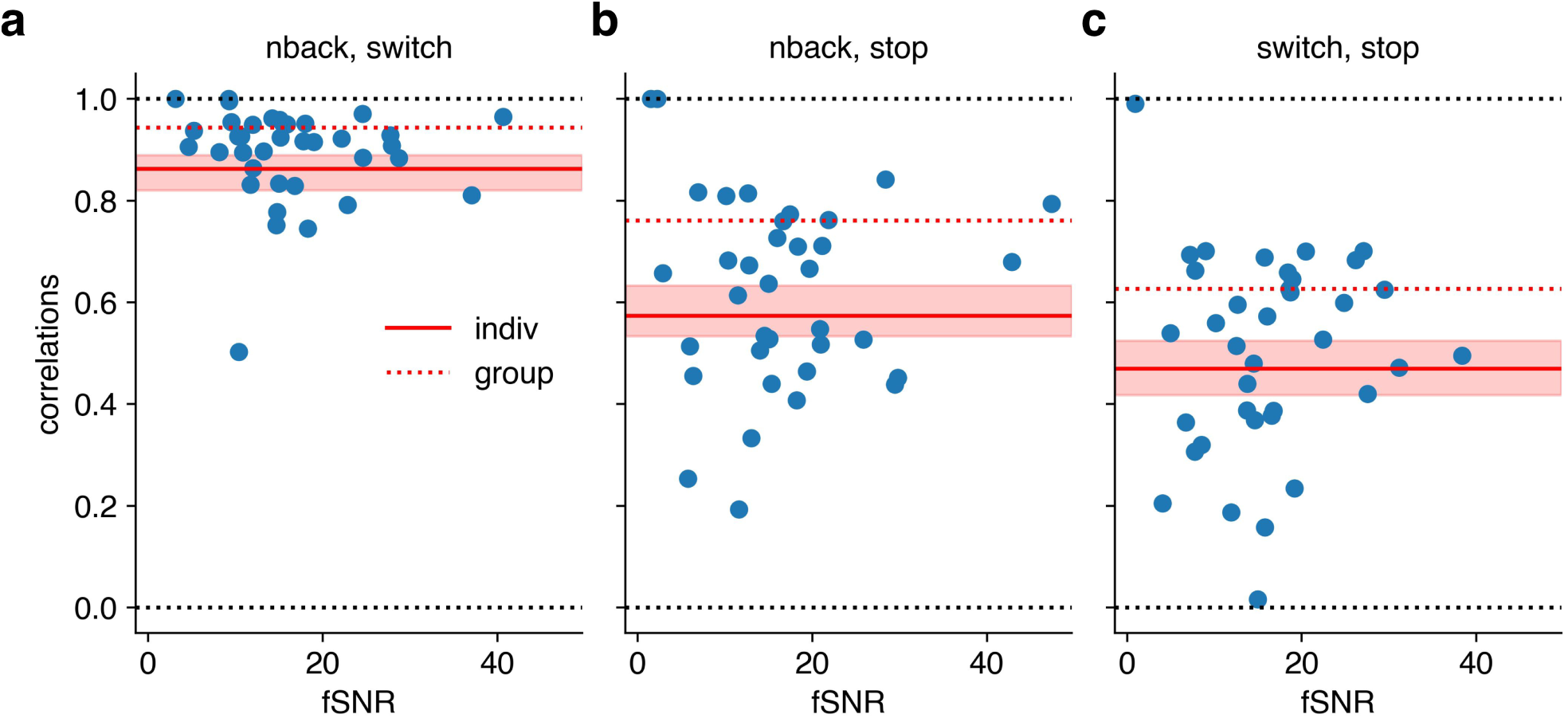
Noise-corrected correlations of activity contrasts across tasks in core MD frontal regions. (**a**) Noise-corrected correlation between activity patterns for the N-back and switch tasks, plotted as a function of functional signal-to-noise ratio (fSNR) (Diedrichsen et al., 2026). Blue dots indicate individual-subject estimates (*n* = 37), obtained by fitting the model separately for each subject. The red solid line shows the group-level estimate obtained by fitting a single model to pooled data across subjects, with shaded regions indicating the 95% confidence interval (200 bootstrap resamples). The dashed red line indicates the correlation estimated from group-averaged data. **(b)** As in **(a)** but for between N-back and stop signal tasks. **(c)** As in **(a)** but for between switch and stop signal tasks.

Consistent with the strong sensitivity of MD regions to task difficulty (Assem et al., 2024, Fedorenko et al., 2013), fSNR was high across subjects. In general, correlations estimated at the individual level were consistently lower than those estimated at the group level, specifically 8.6% lower for the N-back and switch tasks, 24.7% lower between N-back and stop signal tasks, and 25.1% lower between switch and stop signal tasks (see Methods 4.7). Because the correlations were corrected for the amount of measurement noise, this result indicates that the group maps overestimate the degree of functional correspondence between tasks. Averaging across subjects likely inflates apparent similarity by smoothing over idiosyncratic functional organization. As a consequence, MD regions defined at the group level may appear more spatially extensive and functionally homogeneous than they are within individuals.

Despite the reduced overlap, all task pairs remained significantly correlated within individuals (bootstrap tests, 200 bootstrap samples, *p <* 0.001), indicating that MD regions retain a robust domain-general component. The degree of overlap, however, differed across task pairs, with the weakest correspondence observed between the switch and stop-signal tasks (Fig. 4).

## 3. Discussion

The human prefrontal cortex (PFC) supports flexible, higher-order cognition across diverse task demands (Duncan, 2001, 2010, Miller and Cohen, 2001). While a growing body of work has emphasized the presence of functional gradients in PFC (Abdallah et al., 2022, Badre, 2008, O’Reilly, 2010), it remains unclear whether discrete functional boundaries also contribute to its organization (Zhi et al., 2022). Here, we addressed this question using a collection of large-scale multi-task fMRI datasets that densely sample cognitive processes across domains (King et al., 2019, Pinho et al., 2018, Nakai and Nishimoto, 2020, Assem et al., 2024, Barch et al., 2013, Arafat et al., 2026).

### 3.1. Inter-individual variability

Consistent with prior observations in resting-state fMRI (Gratton et al., 2018, Mueller et al., 2013, Gordon et al., 2017a,b), we demonstrate substantial inter-individual variability in PFC activity patterns during task execution across diverse cognitive domains. Such variability could cause group-averaged maps to exhibit gradual functional transitions, even when abrupt boundaries exist within individuals (Dworetsky et al., 2024, Ladwig et al., 2026).

We quantified these observations across six task-based fMRI datasets, offering several advantages over resting-state approaches. Task profiles allow direct assessment of functional correspondence across participants, whereas resting-state alignment must rely on indirectly matching networks by their spatial location (Gordon et al., 2017a, Mueller et al., 2013) or by their connectivity fingerprints (Finn et al., 2015, Xu et al., 2016, Du et al., 2025, Buckner et al., 2013). Our statistical framework additionally separates true inter-individual variability from variability attributable to measurement noise (Diedrichsen et al., 2026, Chen et al., 2025).

We find that inter-individual variability in task-evoked PFC responses is much larger than observed in other brain regions. In PFC, only one third of explainable variance reflects shared group-level topology, which mostly explains differences at coarse spatial scales. Below 8 mm spatial resolution, functional organization is largely idiosyncratic.

### 3.2. Existence of functional boundaries

Given the substantial individual variability in task-evoked responses, we derived an individualized atlas for each subject by combining a group atlas with a limited amount of subject-specific task activation data (Zhi et al., 2025, Nettekoven et al., 2024). Unlike the contiguous parcels in group atlases, individualized parcels were often fragmented and spatially variable across subjects, consistent with recent qualitative observations (Ladwig et al., 2026). These individualized parcellations improved the prediction of brain responses relative to group-based atlases.

Importantly, just showing that parcellation maps have predictive power does not imply the existence of functional boundaries, as functional organization could still vary smoothly due to gradients. To directly test for boundaries, we calculated the distance-controlled boundary coefficient (DCBC) on independent data from the same individual. The DCBC compares the similarity of pairs of vertices within and between regions while controlling for their spatial distance (Zhi et al., 2022). A positive DCBC values indicates that the functional profiles change more abruptly across putative boundaries than within regions (Zhi et al., 2022, King et al., 2019, Nettekoven et al., 2024, Zhi et al., 2025).

Using this approach, we found strong evidence for functional boundaries in individualized parcellations that are not apparent in group atlases. Moreover, more than 97% of the boundaries between parcel pairs corresponded to genuine functional discontinuities, indicating that PFC is organized as a mosaic of functionally distinct regions separated by step-like transitions rather than continuous gradients alone.

Although we matched the number of parcels to the Glasser atlas, the appropriate granularity of PFC organization remains an open question (Zhi et al., 2022). Prior work shows that increasing the number of parcels can improve boundary detection up to a point, after which gains plateau (Zhi et al., 2025). The optimal resolution likely depends on both data availability and analytical goals. While a finer parcellation may capture less prominent subdivisions, using fewer parcels will concentrate on the major functional boundaries.

### 3.3. Task-invariant organization of the prefrontal cortex

Although the functional correlation between different parts of the PFC has been shown to change across tasks (Cole et al., 2013, Braun et al., 2015, Shine et al., 2016, Krienen et al., 2014), our results also indicate substantial stability in its internal functional organization. Individualized parcellations derived from one set of task conditions generalized to independent task sets, with DCBC remaining significantly above zero. This suggests that functional boundaries in PFC are to some degree task-invariant (Zhi et al., 2025, Nettekoven et al., 2026).

This finding is consistent with recent electrophysiology work in non-human primates, showing that topographic maps in lateral PFC, defined by the response similarity of neurons to task conditions, maintain a significant shared spatial correlation structure across tasks, even though they also exhibit some variation from one task to another (Xiang et al., 2025b,a). Together with recent fMRI studies (Nettekoven et al., 2026), these results suggest that although each task co-activates the PFC in a specific way, a clear functional organization becomes visible when neural responses are sampled across a large and diverse enough task set.

### 3.4. Fine-grained spatial scale

Our results show that functional organization in PFC is fine-grained and highly individualized, structured by clear functional boundaries. At spatial scales below 8 mm, organization is almost entirely idiosyncratic, but nonetheless exhibits robust discontinuities between regions. This fragmented and interdigitated structure is consistent with theoretical accounts emphasizing high-dimensional, flexible representations in PFC (Rigotti et al., 2013, Fusi et al., 2016, Mante et al., 2013). Compared to parietal and sensory cortices, PFC exhibits more rapid spatial variation in functional profiles, suggesting a denser packing of distinct computational units, despite strong anatomical connectivity (Goldman-Rakic, 1984, Xu et al., 2022).

Such an organization may support both specialization and integration: adjacent regions may perform related computations while being biased toward different input domains or task demands (Assem et al., 2025, Ladwig et al., 2026, Fedorenko et al., 2013). This architecture enables flexible recombination of functional components, consistent with compositional accounts of cognition (Cole et al., 2011, Ito et al., 2022, Duncan et al., 2017, Yang et al., 2019).

### 3.5. Practical consequences for studying prefrontal organization in humans

Our findings highlight that key aspects of PFC functional organization are only observable at the individual level. Even with advanced alignment techniques, inter-subject variability remains substantial, limiting the interpretability of group-averaged maps (Mueller et al., 2013). This supports a growing shift toward dense, within-subject sampling approaches in human neuroimaging (Gratton and Braga, 2026, Thirion et al., 2021, Poldrack et al., 2015) and individual-level analysis (Dworetsky et al., 2024).

Individualized parcellation offers a promising tool for capturing fine-grained organization. Although resting-state fMRI may be used to derive an individualized parcellation, it fails to capture task-state boundaries, as resting and task states emphasize different aspects of functional organization (Nettekoven et al., 2026, Zhi et al., 2022, Cole et al., 2016). Sampling across multiple tasks provides a more comprehensive basis for identifying functional regions (Nettekoven et al., 2026), and recent work has begun to formalize how such task batteries can be optimized (Arafat et al., 2026).

These findings have important implications for how large-scale systems such as the multiple-demand system (Duncan, 2010, Assem et al., 2020) and frontoparietal networks (Cole et al., 2013, Ji et al., 2019) are interpreted. While typically defined using group-averaged data and described as spatially coherent, our results suggest that, within individuals, these systems may consist of multiple interdigitated sub-regions with distinct functional profiles. Characterizing these systems at the individual level may therefore provide a more precise account of their functional architecture and how domain-general and task-specific processes are integrated (Ladwig et al., 2026, Diedrichsen et al., 2026).

More fundamentally, group analyses emphasize common structure while obscuring individual organization. Consistent with this, cross-task activity correlations in MD cortex were substantially lower within individuals than in group-averaged maps despite high measurement reliability. These observations suggest that domain-general regions may be more appropriately defined by reliable cross-task correspondence within individuals than by consistent activation across subjects. Whether this organizational principle generalizes to broader task spaces remains an important question for future work.

Finally, our conclusions are limited by the spatial resolution of fMRI (∼ 2 mm). Residual structure at this scale suggests that even finer-grained organization remains to be uncovered, potentially accessible with high-field imaging (Yacoub et al., 2008).

### 3.6. Gradients and boundaries as complementary principles

Our results reveal that PFC functional organization is jointly shaped by functional gradients and discrete boundaries. Gradients capture smooth, large-scale variation in functional preference across the cortical surface (Badre, 2008, Abdallah et al., 2022), while boundaries mark abrupt transitions in functional profiles between nearby regions that would otherwise be expected to be similar (Zhi et al., 2022, King et al., 2019, Nettekoven et al., 2024).

One tentative explanation for the co-existence of gradients and boundaries is that adjacent regions occupy similar positions along large-scale functional gradients and therefore perform related computations, while boundaries may separate sub-regions that differ in their input modalities or processing domains (Abdallah et al., 2022, Assem et al., 2025, Ladwig et al., 2026, Fedorenko et al., 2013). Under this account, the gradient reflects a continuous axis of functional preference, and boundaries arise where input or domain specificity shifts abruptly along that axis. This interpretation remains speculative. Future work should test it more directly, for instance, by first identifying domain-general and domain-specialized regions at the individual level, then examining how these map onto the gradient-boundary structure we describe here.

### 3.7. Conclusions

In summary, we show that the human prefrontal cortex exhibits a highly individualized and fine-grained functional organization in which continuous gradients coexist with discrete functional boundaries. While group-level analyses primarily capture coarse gradients (Badre, 2008, Abdallah et al., 2022), individualized approaches reveal a rich mosaic of functionally distinct regions that are stable across tasks.

These findings refine the view of PFC as a spatially homogeneous, domain-general system (Assem et al., 2020, Duncan, 2010) and instead support a more nuanced architecture composed of interdigitated functional sub-regions. This organization provides a potential substrate for the flexibility and compositionality that characterize higher-order cognition (Yang et al., 2019, Duncan et al., 2017, Duncan, 2001, Rigotti et al., 2013).

## 4. Methods

### 4.1. Datasets and data organization

We used six task-based fMRI datasets (see Supplementary Table 1). All studies were approved by the ethics committee and review board at respective institutes. Each of the first four datasets leveraged a broad battery of tasks that tap into multiple processing domains, including cognitive, social, affective, motor and perceptual functions: (1) The *multi-domain task battery* (MDTB) dataset (King et al., 2019); (2) the *Nakai & Nishimoto* dataset (Nakai and Nishimoto, 2020); (3) the *individual brain charting* (IBC) dataset (Pinho et al., 2018); (4) the task data for the *unrelated 100* subjects in the *Human Connectome Project* (HCP) S1200 2017 release (Barch et al., 2013). In addition, we also included two datasets for a better description of high-order execution and language: (5) the *multiple-demand* dataset (Assem et al., 2024); (6) the *language task battery* dataset (Arafat et al., 2026).

The task-based datasets were preprocessed as described in Zhi et al. (2025). The preprocessing pipeline included correction for spatial distortion and head motion, registration to the structural data, cortical surface mapping and functional artifact removal. No smoothing or group normalization was applied at this stage (Nettekoven et al., 2024). The beta weights from the first-level GLM were univariately pre-whitened by dividing them by the square root of the residual mean-square image Zhi et al. (2022). Using a unified framework (available at github.com/DiedrichsenLab/Functional_Fusion), the data were then extracted in fs32k atlas space.

### 4.2. Decomposing the variance into group, subject and noise components

The data for each participant were partitioned into two halves, where each half contained the same task conditions. The data *y* can be represented as a 4-dimensional tensor of beta weights with shape *N* × *R* × *K* × *P*, where *N* is the number of participants, *R* the number of partitions, *K* the number of task conditions and *P* the number of vertices. We decomposed the data into three components (1) a group signal *g* that is consistent across participants and partitions; (2) a signal *s* that is unique to a subject and consistent across partitions; (3) noise *ε* that fluctuates across partitions (Seghier and Price, 2018, Ejaz et al., 2015). The three components are orthogonal to each other, and fully capture the total variability in the data. More specifically, for participant *i*, partition *j*, *y_ij_* = *g* + *s_i_* + *ε_ij_*.

The variances for *g*, *s* and *ε* can be derived as follows:

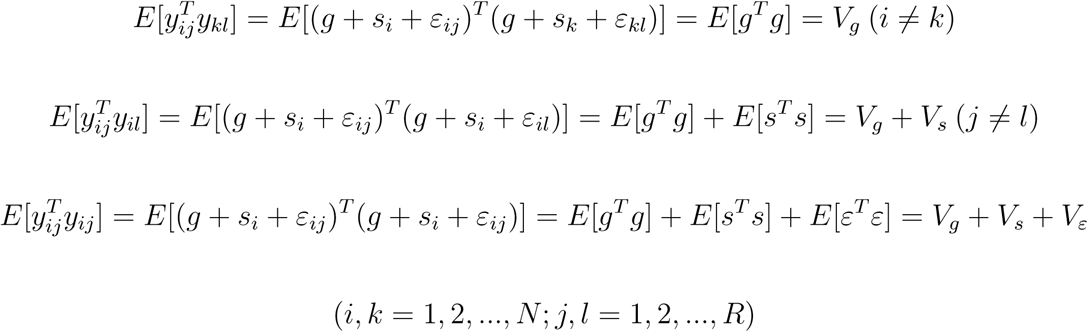

We estimated the mathematical expectancies using all possible pairs of measurements across participants and partitions. The variance decomposition can be done vertex-wise (*K* × 1) or for the multi-voxel response patterns (*K* × *P*).

If rest was not included as a condition in a dataset, 0 was used as an estimate for activation during rest (King et al., 2019). The beta weights for each voxel were mean-centered across conditions before variance decomposition.

Variance decomposition in Fig. 1a was done vertex-wise. Results were smoothed using a 2-dimensional (2D) Gaussian kernel with full-width-at-half-maximum (FWHM) of 4 mm for each dataset before averaging across datasets. Because cross-validated variance estimates can be noisy when signal quality is low, occasional estimates fell outside the theoretically plausible range. To prevent such extreme values from spreading during spatial smoothing, variance estimates were clipped to the range [−0.5, 1.5] prior to smoothing, with values outside this interval set to the corresponding boundary value.

Variance decomposition in Fig. 1b was done for the multi-voxel response patterns. We estimated *V_g_*, *V_s_*, and *V_ε_* for each subject. Results were summarized across subjects and datasets.

Variance decomposition in Fig. 1c was done globally. Results were summarized across datasets.

### 4.3. Dividing neural activity into non-overlapping spatial frequency bands

To assess inter-individual variability at different spatial scales, we split the data into multiple non-overlapping spatial frequency bands (≥ 10 mm/cycle, 8 – 10 mm/cycle, 6 – 8 mm/cycle, 4 – 6 mm/cycle, and 2 – 4 mm/cycle) by applying an iterative surface-smoothing and subtraction procedure. More specifically, to get the signals that are coarser than 10 mm/cycle, for each participant, partition, and task condition, we smooth the original data using a 2D Gaussian kernel with FWHM of 10 mm. We subtract out the 10 mm smoothed signal from the original signal, and smooth the residuals with a 2D Gaussian kernel of 8 mm FWHM to get the signals that are coarser than 8 mm/cycle but finer than 10 mm/cycle. Repeating this step, we can get the signals for other frequency bands.

### 4.4. Distance-controlled boundary coefficient (DCBC)

Existing metrics for evaluating parcellation, such as the global Homogeneity and Silhouette coefficient, do not take into account the intrinsic smoothness of functional data, where nearby voxels have similar response profiles (Gordon et al., 2016, Rousseeuw, 1987, Craddock et al., 2012). More specifically, global Homogeneity measure would compute the average response profile correlations across all pairs of voxels in a parcel; Silhouette coefficient would compute the difference of tuning profiles for between-parcel and within-parcel using all pairs of voxels. Due to the intrinsic smoothness in the functional data, these global measures are biased by the resolution of parcels or by the fact that within-parcel voxel pairs tend to be closer than between-parcel voxel pairs (Zhi et al., 2022).

It is therefore critical to only compare the within– and between-parcel functional profile similarities using voxel pairs separated by the same spatial distance. This is exactly what DCBC does. DCBC first computes the correlations between the functional profiles of vertex pairs that are assigned within– and between-parcels at certain spatial bin, with cross-validation across partitions. The cross-validated correlation is estimated using the following formula:

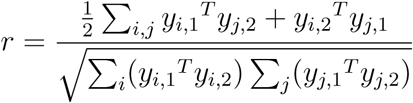

where *y_i,_*_1_ represents the response profile of vertex *i* in partition 1. The distances among vertices on the cortical surface are calculated using geodesic distance, approximated using Dijkstra algorithm (Dijkstra, 1959). Distances were binned every 5 mm up to 35 mm. To get a simple evaluation criterion, we then integrate the difference of within– and between-parcel functional profile correlations across spatial bins via weighted averaging. The weight for the *i*-th bin is defined follows:

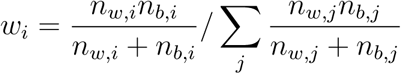

where *n_w,i_*refers to the number of within-parcel vertex pairs in the *i*-th bin, while *n_b,i_* refers to the number of between-parcel vertex pairs in the *i*-th bin.

Spatial distances among vertices are approximated using Dijkstra’s algorithm to estimate the shortest paths between each pair of vertices on each individual’s cortical surface (Dijkstra, 1959). We compute the pairwise distances using the mid-cortical layer for each individual, which is the average of the pial and white-gray matter surface, then average the distances across individuals. This resulted in a matrix that indicates the average cortical distance between nearby brain locations for the atlas brain surface (Zhi et al., 2022).

The DCBC values (integrated across bins) are independent of the resolution of the parcels and be compared directly against 0. Random parcellation or smoothly varying gradients would yield DCBC of 0, indicating no real boundaries between parcels.

More details on DCBC can be found at (Zhi et al., 2022).

### 4.5. Individualized brain parcellation

We applied a hierarchical Bayesian parcellation (HBP) framework (Zhi et al., 2025) to derive individualized brain parcellations for each MDTB subject by combining a group-level atlas with limited subject-specific task data. In this framework, the individualized parcellation *U^s^* is treated as a latent variable and estimated by maximizing the data log-likelihood using an Expectation–Maximization (EM) algorithm (Bishop, 2009).

The model incorporates a probabilistic group atlas as a prior, specifying the probability that each brain location belongs to each of *K* parcels. To construct this prior, the Glasser atlas was first converted into a one-hot encoding of parcel assignments and then transformed into a probabilistic atlas using a softmax function with strength parameter *θ* = 20.

At each EM iteration, the E-step estimates the posterior probability that brain location *i* belongs to parcel *k* for a given subject, combining the likelihood of the individual data with the group prior. In the M-step, model parameters are updated to maximize the expected log-likelihood given these posterior assignments. The data likelihood was modeled using a mixture of von Mises–Fisher (vMF) distributions, which has been shown to effectively capture functional profiles across brain regions (Zhi et al., 2025, Banerjee et al., 2005, Lashkari et al., 2010, Ryali et al., 2013, Schaefer et al., 2018, Thomas Yeo et al., 2011). The parcel-specific mean response vectors (*V*) and the concentration parameter (*κ*) were iteratively updated during optimization. The EM procedure was repeated until convergence of the log-likelihood. A final E-step was then performed, and each brain location was assigned to the parcel with the highest posterior probability, yielding an individualized parcellation.

Further methodological details are provided in (Zhi et al., 2025).

### 4.6. Spatial autocorrelation function

To investigate the smoothness and spatial scale of the functional organization, we first compute a cross-validated covariance matrix and a cross-validated variance matrix, which will be used to compute cross-validated spatial autocorrelation function.

For computing a spatial autocorrelation curve, as in Fig. 3a, we estimate the average covariance divided by the average variance across all pairs of vertices spaced at a certain distance bin for each subject. This provides a global estimate of the spatial scale.

The distance estimates are the same as in DCBC. At distance 0, cross-validated covariances equal to cross-validated variances, yielding a cross-validated spatial autocorrelation of 1.

To estimate the smoothness at each brain location and create a map as in Fig. 3b, for each row of the two matrices, we estimate the average covariance and average variance within a spatial distance bin. The ratio of which provides an estimate of how the response profile of a particular vertex is correlated with all the other cortices at the given distance. We then fit the spatial autocorrelation function for each vertex with an exponential function:

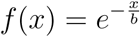

where x stands for the distance bin in millimeter and *b* is the parameter to be estimated. The fitted parameter can be used to estimate the FWHM of the exponential function as an indicator for local smoothness:

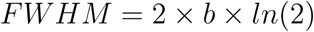

We do so for each subject and then average the FWHMs for each vertex across subjects for visualization.

### 4.7. Noise-corrected correlations of activity contrasts across tasks in core MD frontal regions

We used a multiple-demand fMRI dataset (Assem et al., 2024) to examine the implications of our findings for the multiple-demand (MD) system (Duncan, 2010, Assem et al., 2020). In this dataset, each subject performed three tasks (N-back, task switching, and stop signal), each with two difficulty levels (hard vs. easy). The difficulty contrast (hard–easy) provides a robust functional signature of the MD system (Fedorenko et al., 2013, Duncan, 2010, Assem et al., 2024). Additional details are provided in Methods 4.1, Supplementary Information 1, and Assem et al. (2024).

To quantify the similarity of activity patterns across tasks, we used a noise-corrected correlation metric that estimates the correlation between true activity patterns while accounting for measurement noise (Diedrichsen et al., 2026). In this framework, signal and noise variance–covariance structures are modeled separately. Under the assumptions of normally distributed signal and noise and independence across voxels, maximum-likelihood estimates of the true correlation, signal variance, and noise variance can be obtained (Diedrichsen et al., 2026). The functional signal-to-noise ratio (fSNR) was defined as the ratio of signal variance to noise variance.

We estimated noise-corrected correlations of activity contrasts across task pairs using three approaches: (i) individual-level estimates obtained by fitting the model separately for each subject, (ii) a group-level estimate obtained by fitting a single model to data pooled across subjects, and (iii) estimates derived from group-averaged data. This comparison allowed us to dissociate the effects of inter-individual variability and group averaging on correlation estimates. For statistical inference, we performed a group bootstrap analysis by resampling participants with replacement (200 bootstrap samples). For each resampled dataset, group-level correlation estimates were recomputed to derive confidence intervals (Diedrichsen et al., 2026).

All analyses were restricted to core MD frontal regions as defined in (Assem et al., 2020).

## 5. Data availability

The MDTB dataset is publicly available at https://openneuro.org/datasets/ds002105/versions/1.1.0.

The Nakai and Nishimoto dataset is publicly available at https://osf.io/ea2jc/ and https://openneuro.org/datasets/ds002306.

The HCP task fMRI data used in this study are publicly available from the Human Connectome Project (https://www.humanconnectome.org).

The Individual Brain Charting dataset is publicly available at https://project.inria.fr/IBC/.

The Multiple-Demand dataset is publicly available at https://balsa.wustl.edu/study/0qk6K.

The Language dataset is publicly available at https://openneuro.org/datasets/ds007276/versions/1.1.0.

## 6. Code availability

Code used for analysis is available at GitHub github.com/jkderrick028/PFC_parcellation. git.

## 7. Author contributions

JDX, MM, JD conceptualized the work. DZ, JD designed the Hierarchical Bayesian Parcellation framework and the DCBC metric. BA, DZ, CN, JD maintained the Functional Fusion framework and related datasets. BA acquired the Language dataset. JD developed the framework for estimating noise-corrected correlations. JDX analyzed the data and wrote the manuscript. MM, JD provided analysis advice. JDX, MM, JD edited the manuscript.

## 8. Acknowledgments

This work was supported by the Natural Sciences and Engineering Research Council of Canada (NSERC) Discovery Grant Program (MM, JD) and the Mitacs Globalink Graduate Fellowship (JDX). Additional support was provided by BrainsCAN at University of Western Ontario through the Canada First Research Excellence Fund (CFREF).

DZ is supported by the National Institute of Mental Health (NIMH) under grant R01MH130899. CN is supported by a Wellcome Trust Early Career Award (306553/Z/23/Z) and a Junior Research Fellowship Grant from Linacre College, University of Oxford.

We thank Ana Lúısa Pinho for helpful discussion and feedback.

## 9. Disclosures

The authors declare no competing interests.

## 10. Supplementary Information

### 10.1. Multi-task fMRI datasets

**Table 1:**
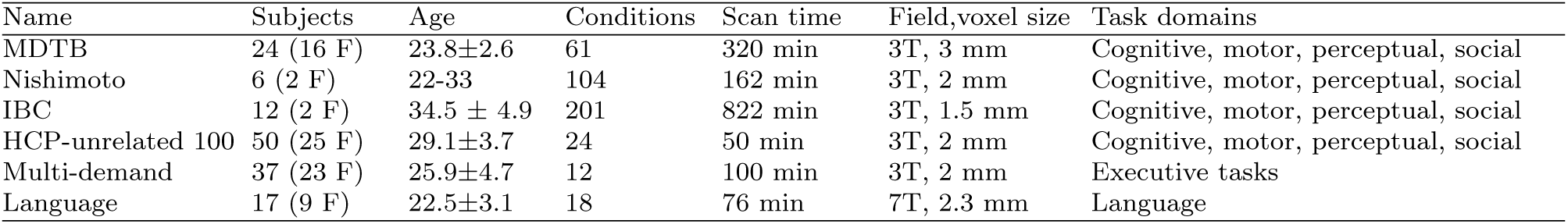
FMRI datasets analyzed in the current paper.

| Name | Subjects | Age | Conditions | Scan time | Field, voxel size | Task domains |
| --- | --- | --- | --- | --- | --- | --- |
| MDTB | 24 (16 F) | 23.8±2.6 | 61 | 320 min | 3T, 3 mm | Cognitive, motor, perceptual, social |
| Nishimoto | 6 (2 F) | 22-33 | 104 | 162 min | 3T, 2 mm | Cognitive, motor, perceptual, social |
| IBC | 12 (2 F) | 34.5 ± 4.9 | 201 | 822 min | 3T, 1.5 mm | Cognitive, motor, perceptual, social |
| HCP-unrelated 100 | 50 (25 F) | 29.1±3.7 | 24 | 50 min | 3T, 2 mm | Cognitive, motor, perceptual, social |
| Multi-demand | 37 (23 F) | 25.9±4.7 | 12 | 100 min | 3T, 2 mm | Executive tasks |
| Language | 17 (9 F) | 22.5±3.1 | 18 | 76 min | 7T, 2.3 mm | Language |

In the table, F stands for female. Age is reported in years (mean ± standard deviation, or range). Scan time is reported in minutes per subject.

### 10.2. Brain regions of interest table

We define brain regions of interest using a subset of Glasser parcels. For PFC, we use the same set of parcels as in (Donahue et al., 2018).

**Table 2:**
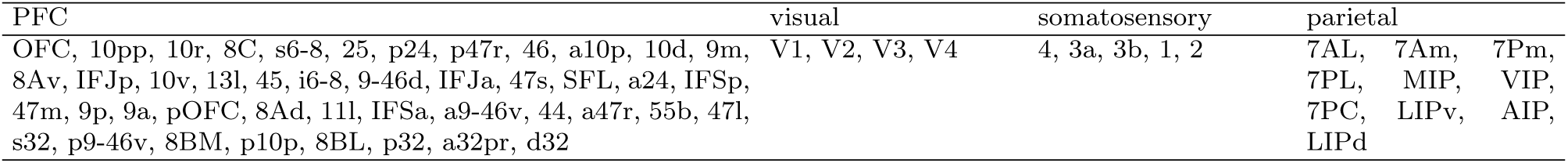
Brain regions of interest.

### 10.3. DCBC on group and individualized Schaefer atlas

**Supplementary Figure 1:**
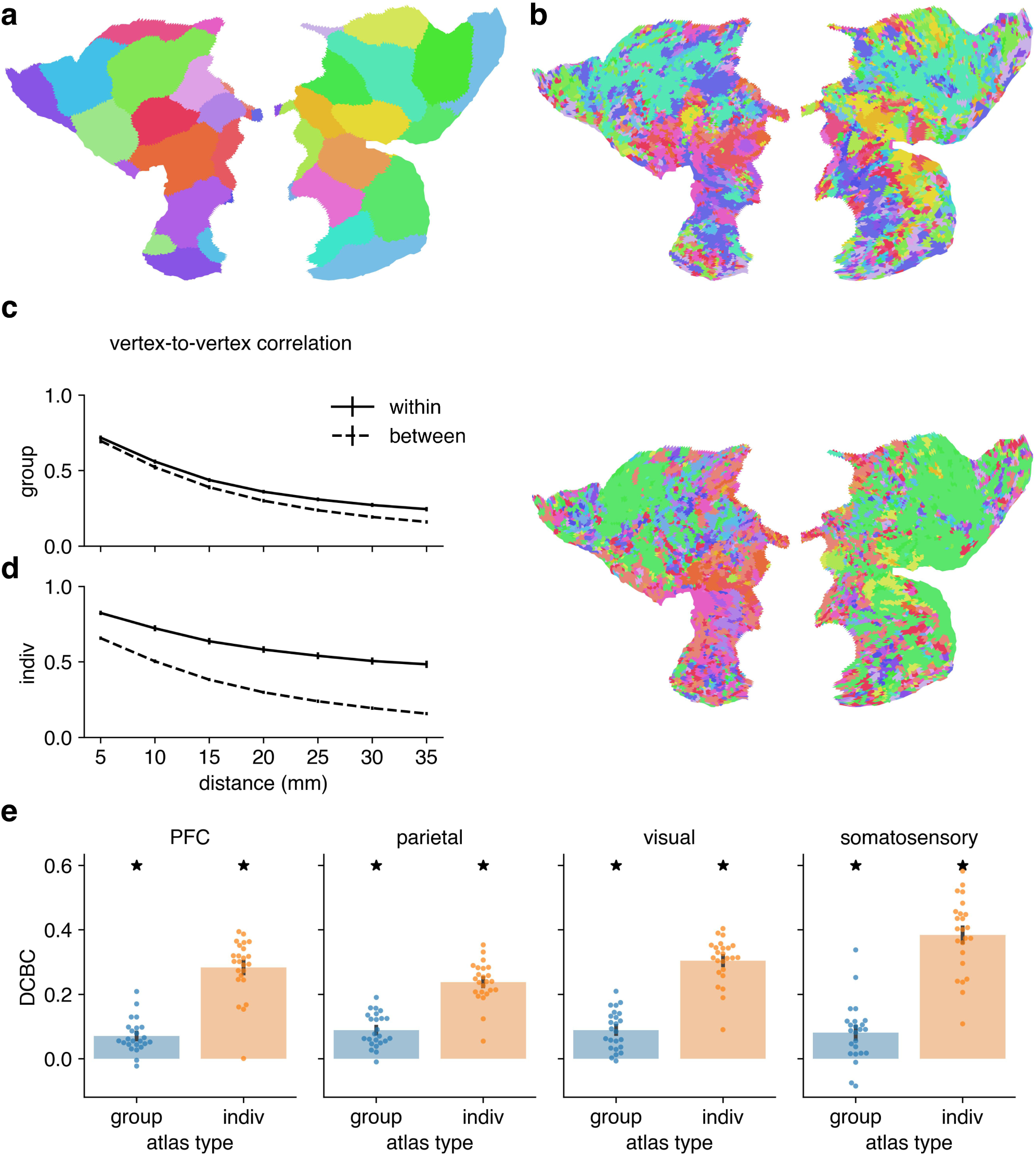
DCBC Schaefer. **(a)** Schaefer group atlas for PFC (Schaefer et al., 2018). **(b)** Individualized Schaefer atlas for the same two example subjects as in Fig. 2b. **(c)** Average cross-validated correlation as a function of spatial distance for functional parcel boundaries for the Schaefer group atlas. The DCBC is defined as the difference in correlation (within – between) within each distance bin. The error bars show standard error across participants (*n* = 24). **(d)** As in **(c)**, but for the individualized Schaefer atlas. **(e)** Average DCBC for the Schaefer group (blue) and individualized (orange) atlases across subjects (*n* = 24). Error bars show standard error across subjects. Asterisks indicate that DCBC is significantly higher than 0 (one-sided t-test, *p <* 0.05, uncorrected). For each ROI, DCBC on the individualized atlas is significantly higher than the group (two-sided t-test, *p <* 0.05, uncorrected). Significance not shown for simplicity.

### 10.4. Group and individualized Glasser atlases for all the other ROIs

**Supplementary Figure 2:**
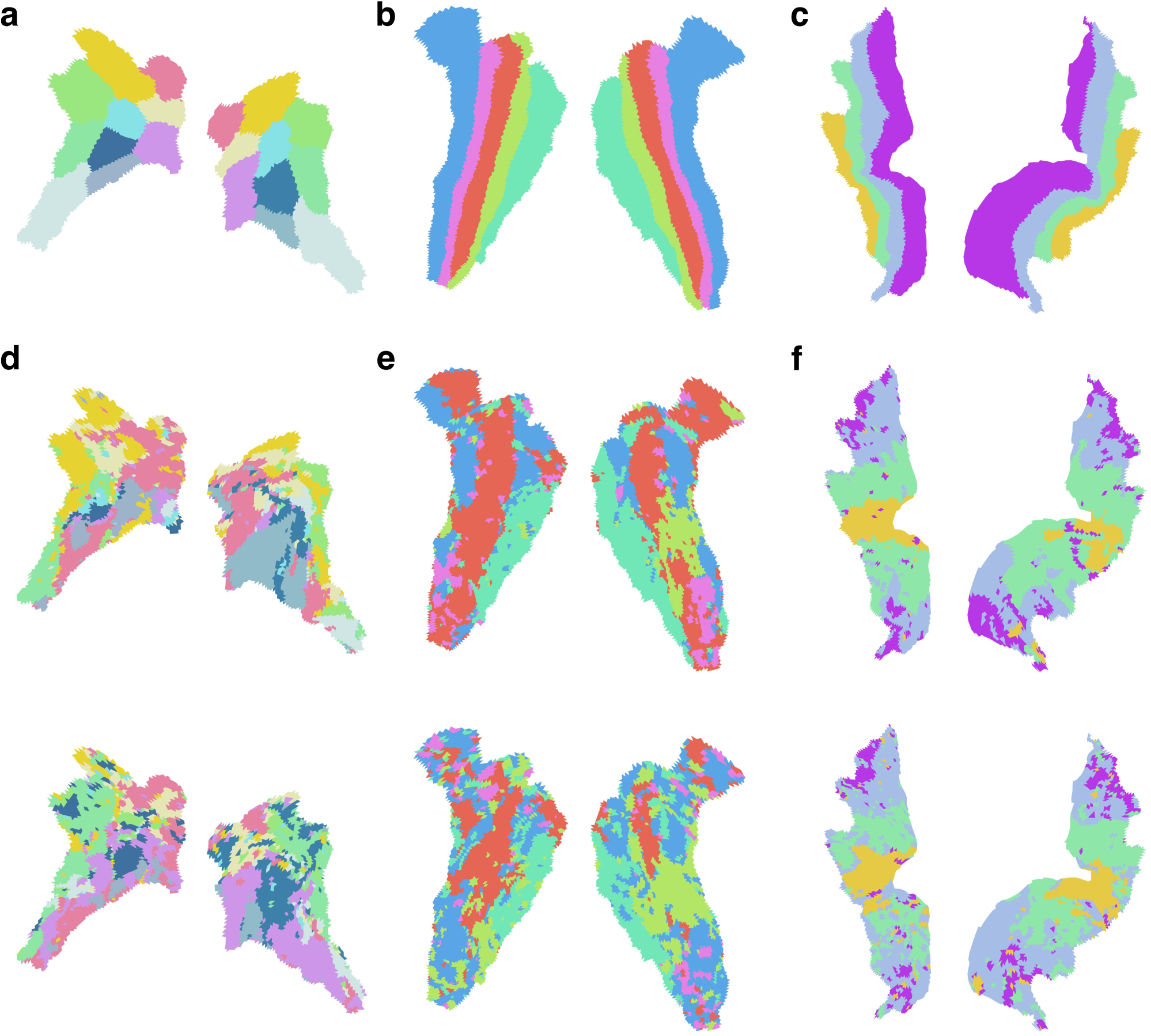
Group and individualized Glasser atlases for the other ROIs. **(a)**-**(c)** Glasser group atlas for for parietal, somatosensory and visual cortices. **(d)**-**(f)** Individualized Glasser atlas in example subjects (same cohort as in Fig. 2b).

Given the substantial inter-subject variability in neural activity in PFC, the Glasser group atlas (Glasser et al., 2016), which is based averaging functional data across subjects, is expected to perform sub-optimally in terms of revealing functional boundaries. However, in brain areas that have predominantly group topology, such as the visual regions (Fig. 1a,b), the Glasser group atlas also fails to reveal functional boundaries (Fig. 2e). This is because that in the Glasser group atlas, the boundaries are identified according to cytoarchitectonic regions (V1, V2, V3), and are vertically oriented. However, a parcellation based on functional data, such as the Schaefer atlas (Schaefer et al., 2018) and the individualized Glasser atlas, separate areas associated with foveal and peripheral vision and are horizontally oriented (Supplementary Fig. 2b). The points out that functional organization does not always align with structural organization (Straathof et al., 2019).

### 10.5. Group and individualized Schaefer atlases for all the other ROIs

**Supplementary Figure 3:**
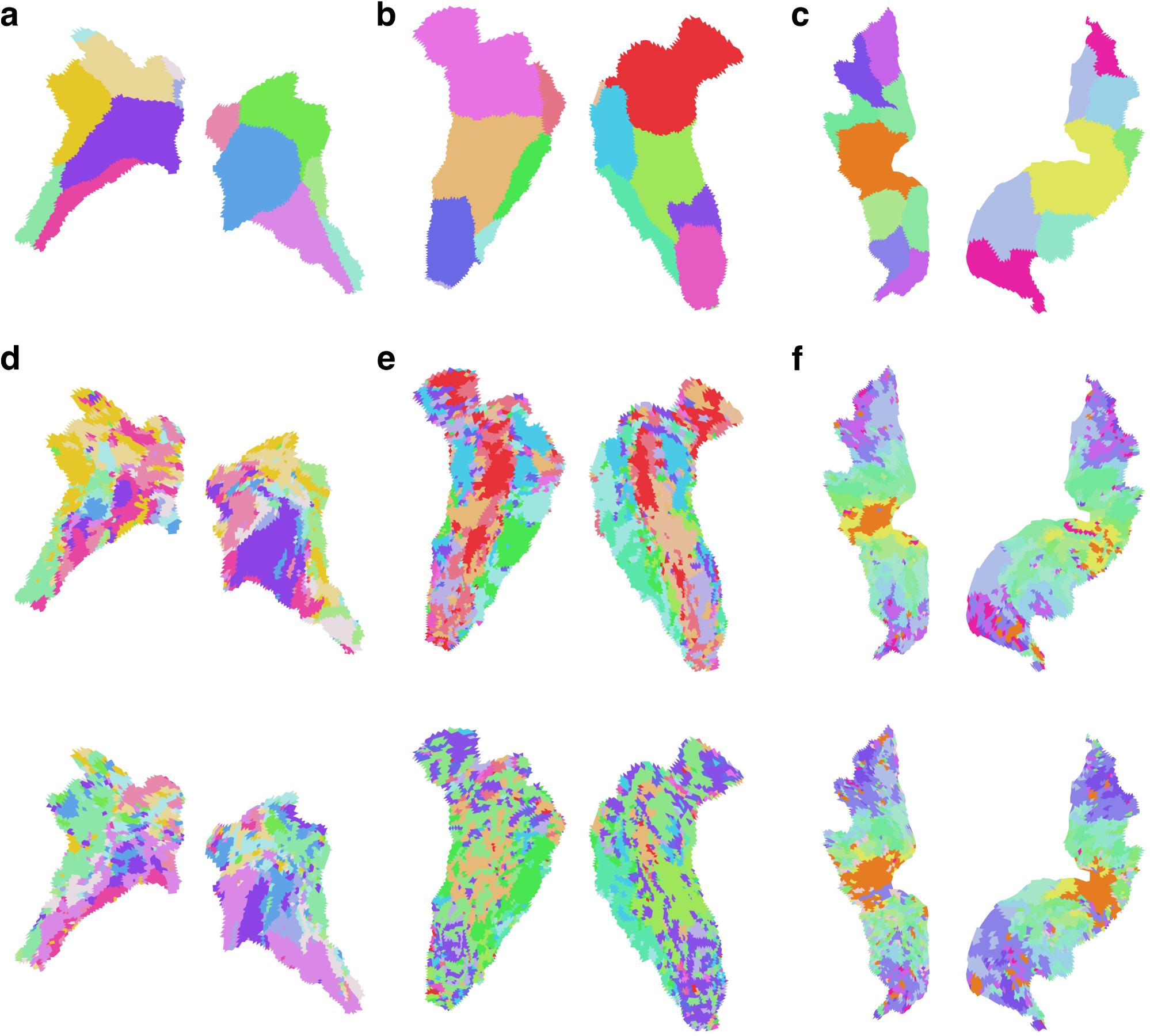
Group and individualized Schaefer atlases for the other ROIs. **(a)**-**(c)** Schaefer group atlas for for parietal, somatosensory and visual cortices. **(d)**-**(f)** Individualized Schaefer atlas in example subjects (same cohort as in Fig. 2b).

### 10.6. Spatial scale of functional organization across datasets

**Supplementary Figure 4:**
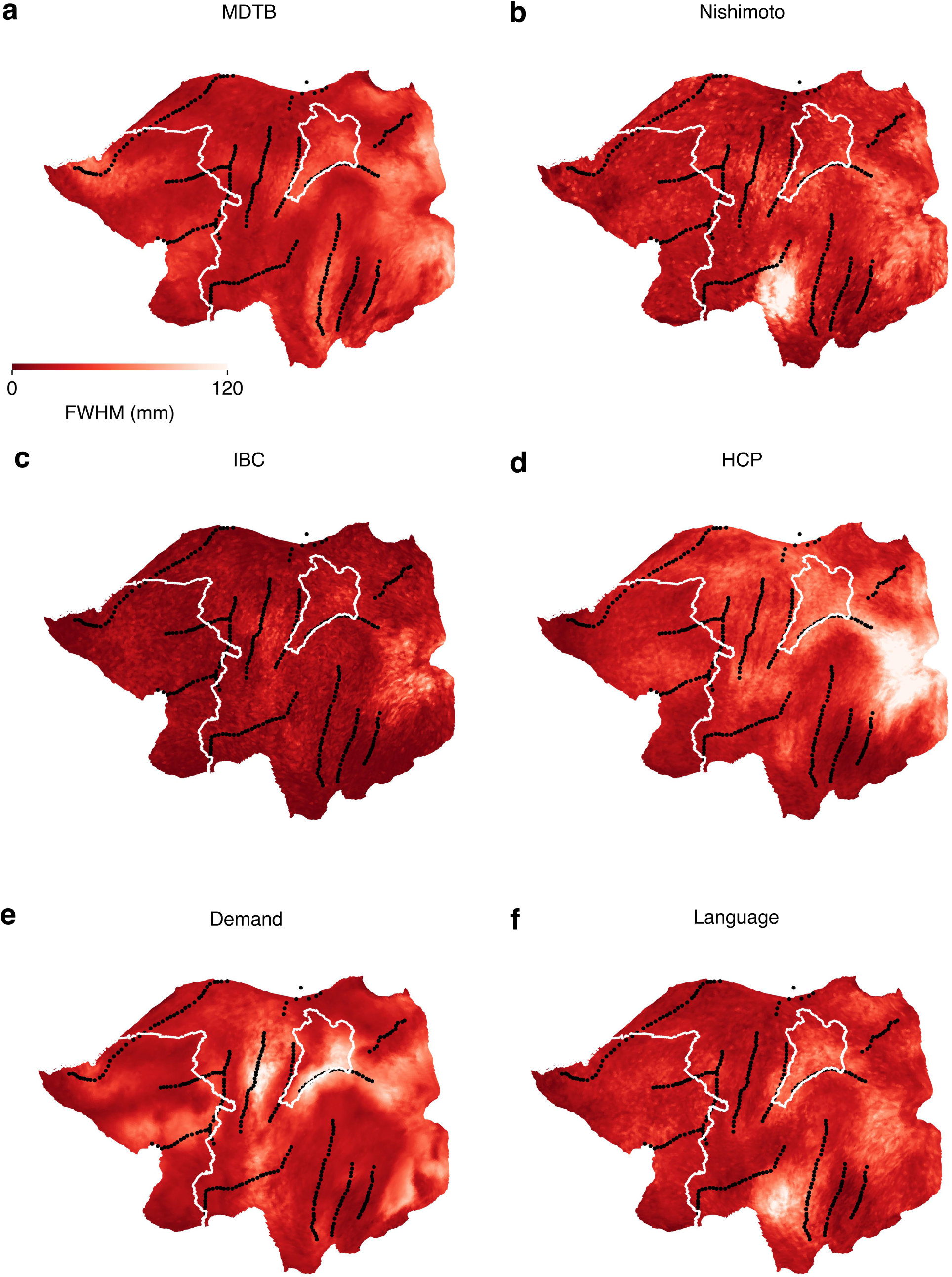
The spatial scale of functional organization in each dataset. **(a)** Full-width-at-half-maximum (FWHM) of the spatial autocorrelation function for each vertex, averaged across subjects in the MDTB dataset. **(b)-(f)**As in **(a)**but for Nishimoto, IBC, HCP, Demand and Language datasets respectively.

## Notes

### Competing Interest Statement

The authors have declared no competing interest.

