## supplementary_materials for "Individualized parcellation reveals functional boundaries in human prefrontal cortex"

### 10. Supplementary Information

#### *10.1. Multi-task fMRI datasets*

| Name | Subjects | Age | Conditions | Scan time | Field,voxel size | Task domains |
|---|---|---|---|---|---|---|
| MDTB | 24 (16 F) | 23.8±2.6 | 61 | 320 min | 3T, 3 mm | Cognitive, motor, perceptual, social |
| Nishimoto | 6 (2 F) | 22-33 | 104 | 162 min | 3T, 2 mm | Cognitive, motor, perceptual, social |
| IBC | 12 (2 F) | 34.5 ± 4.9 | 201 | 822 min | 3T, 1.5 mm | Cognitive, motor, perceptual, social |
| HCP-unrelated 100 | 50 (25 F) | 29.1±3.7 | 24 | 50 min | 3T, 2 mm | Cognitive, motor, perceptual, social |
| Multi-demand | 37 (23 F) | 25.9±4.7 | 12 | 100 min | 3T, 2 mm | Executive tasks |
| Language | 17 (9 F) | 22.5±3.1 | 18 | 76 min | 7T, 2.3 mm | Language |

**Table 1:** FMRI datasets analyzed in the current paper

In the table, F stands for female. Age is reported in years (mean ± standard deviation, or range). Scan time is reported in minutes per subject.

#### *10.2. Brain regions of interest table*

We define brain regions of interest using a subset of Glasser parcels. For PFC, we use the same set of parcels as in (Donahue et al., 2018).

| PFC | visual | somatosensory | parietal |
|---|---|---|---|
| OFC, 10pp, 10r, 8C, s6-8, 25, p24, p47r, 46, a10p, 10d, 9m, 8Av, IFJp, 10v, 13l, 45, i6-8, 9-46d, IFJa, 47s, SFL, a24, IFSp, 47m, 9p, 9a, pOFC, 8Ad, 11l, IFSa, a9-46v, 44, a47r, 55b, 47l, s32, p9-46v, 8BM, p10p, 8BL, p32, a32pr, d32 | V1, V2, V3, V4 | 4, 3a, 3b, 1, 2 | 7AL, 7Am, 7Pm, 7PL, MIP, VIP, 7PC, LIPv, AIP, LIPd |

**Table 2:** Brain regions of interest

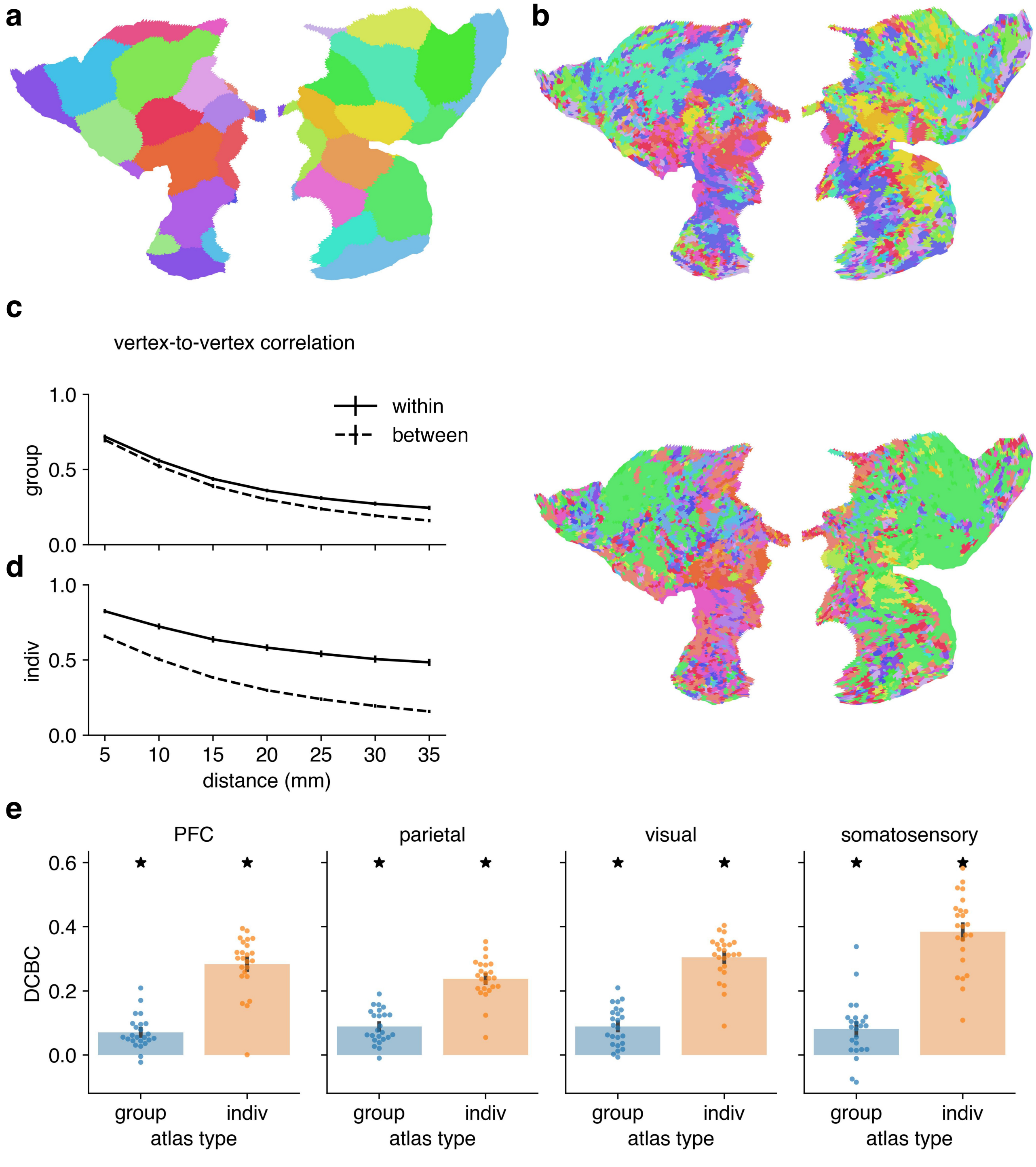


**Supplementary Figure 1: DCBC Schaefer.** **(a)** Schaefer group atlas for PFC (Schaefer et al., 2018). **(b)** Individualized Schaefer atlas for the same two example subjects as in Fig. 2b. **(c)** Average cross-validated correlation as a function of spatial distance for functional parcel boundaries for the Schaefer group atlas. The DCBC is defined as the difference in correlation (within - between) within each distance bin. The error bars show standard error across participants ($n = 24$). **(d)** As in **(c)**, but for the individualized Schaefer atlas. **(e)** Average DCBC for the Schaefer group (blue) and individualized (orange) atlases across subjects ($n = 24$). Error bars show standard error across subjects. Asterisks indicate that DCBC is significantly higher than 0 (one-sided t-test, $p < 0.05$, uncorrected). For each ROI, DCBC on the individualized atlas is significantly higher than the group (two-sided t-test, $p < 0.05$, uncorrected). Significance not shown for simplicity.

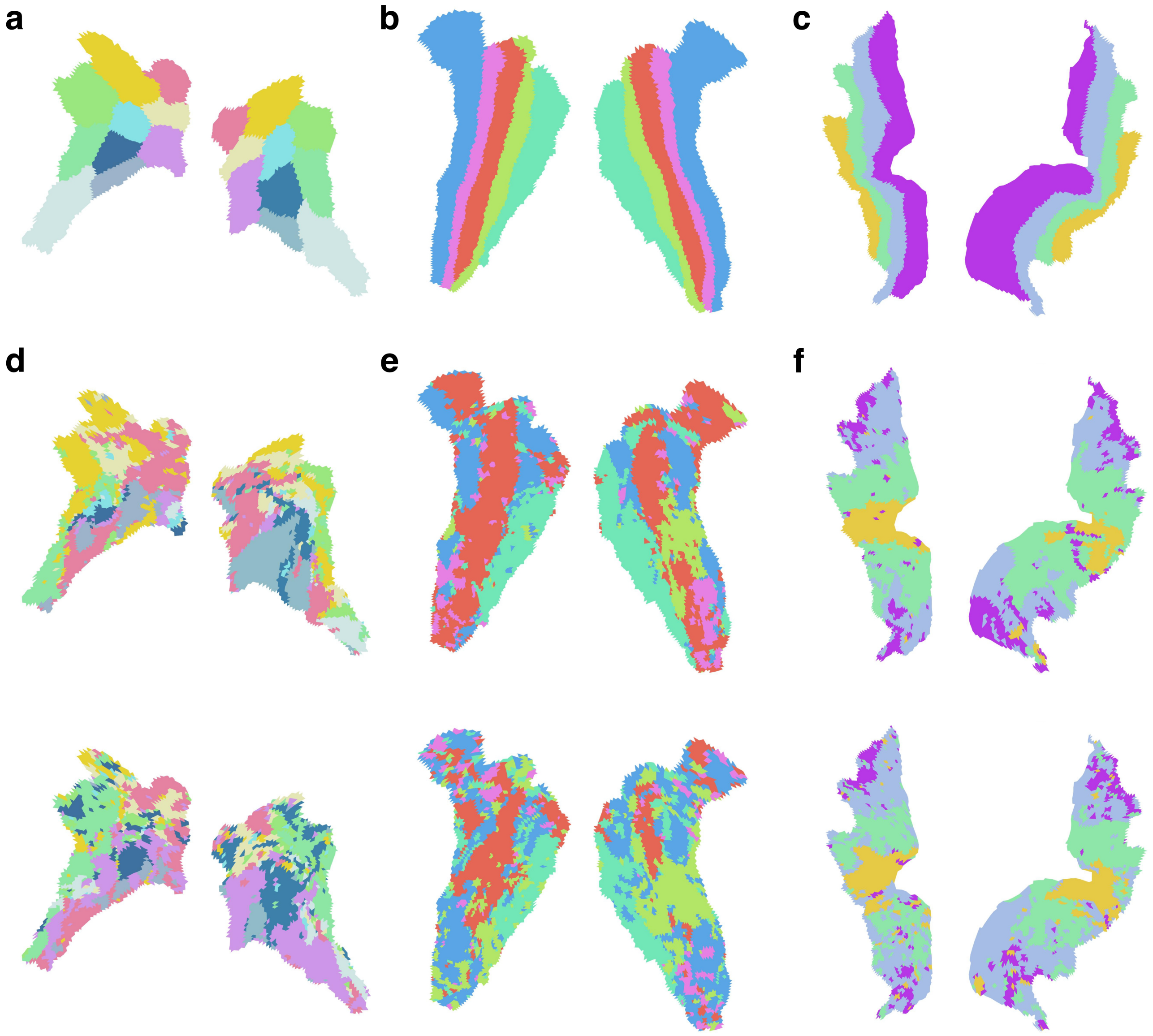


**Supplementary Figure 2: Group and individualized Glasser atlases for the other ROIs. (a)-(c)** Glasser group atlas for for parietal, somatosensory and visual cortices. **(d)-(f)** Individualized Glasser atlas in example subjects (same cohort as in Fig. 2b).

Given the substantial inter-subject variability in neural activity in PFC, the Glasser group atlas (Glasser et al., 2016), which is based averaging functional data across subjects, is expected to perform sub-optimally in terms of revealing functional boundaries. However, in brain areas that have predominantly group topology, such as the visual regions (Fig. 1a,b), the Glasser group atlas also fails to reveal functional boundaries (Fig. 2e). This is because that in the Glasser group atlas, the boundaries are identified according to cytoarchitectonic regions (V1, V2, V3), and are vertically oriented. However, a parcellation based on functional data, such as the Schaefer atlas (Schaefer et al., 2018) and the individualized Glasser atlas, separate areas associated with foveal and peripheral vision and are horizontally oriented (Supplementary Fig. 2b). The points out that functional organization does not always align with structural organization (Straathof et al., 2019).

*10.5. Group and individualized Schaefer atlases for all the other ROIs*

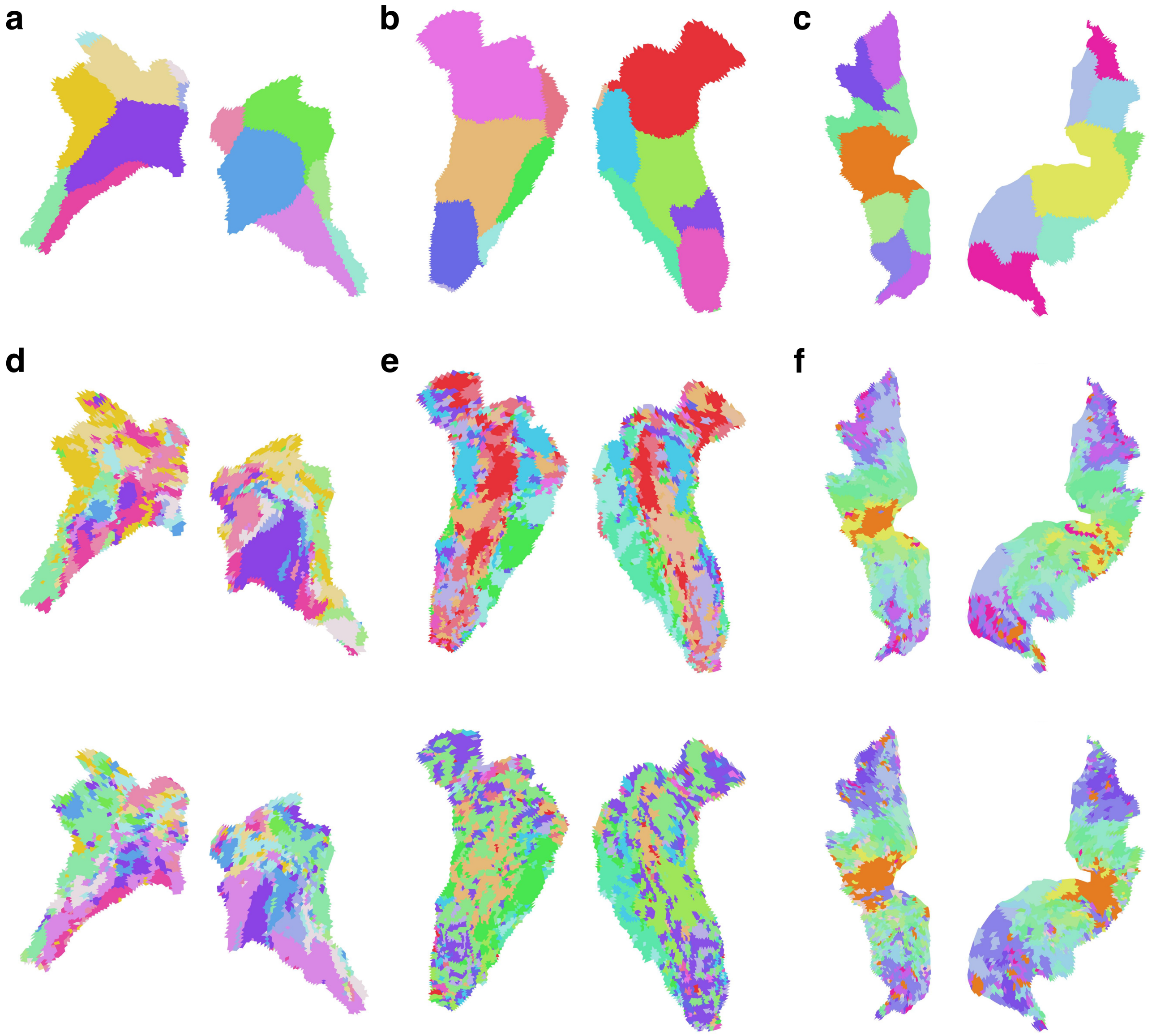


**Supplementary Figure 3: Group and individualized Schaefer atlases for the other ROIs.** **(a)-(c)** Schaefer group atlas for for parietal, somatosensory and visual cortices. **(d)-(f)** Individualized Schaefer atlas in example subjects (same cohort as in Fig. 2b).

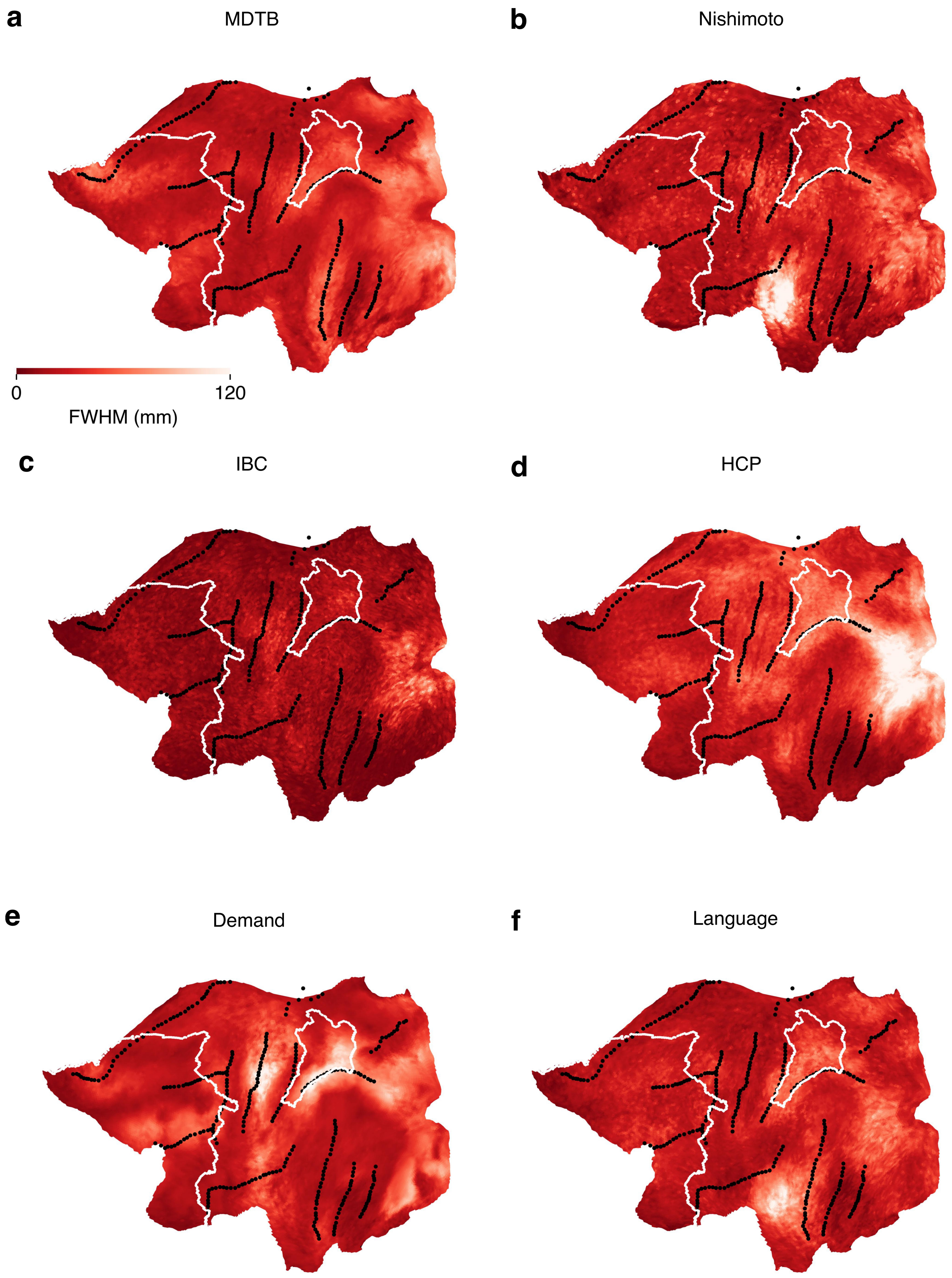


**Supplementary Figure 4: The spatial scale of functional organization in each dataset. (a)** Full-width-at-half-maximum (FWHM) of the spatial autocorrelation function for each vertex, averaged across subjects in the MDTB dataset. **(b)-(f)** As in **(a)** but for Nishimoto, IBC, HCP, Demand and Language datasets respectively.

R. A. Poldrack, T. O. Laumann, O. Koyejo, B. Gregory, A. Hover, M.-Y. Chen, K. J. Gorgolewski, J. Luci, S. J. Joo, R. L. Boyd, S. Hunicke-Smith, Z. B. Simpson, T. Caven, V. Sochat, J. M. Shine, E. Gordon, A. Z. Snyder, B. Adeyemo, S. E. Petersen, D. C. Glahn, D. Reese Mckay, J. E. Curran, H. H. H. Göring, M. A. Carless, J. Blangero,

R. Dougherty, A. Leemans, D. A. Handwerker, L. Frick, E. M. Marcotte, and J. A. Mumford. Long-term neural and physiological phenotyping of a single human. *Nature Communications*, 6(1):8885, Dec. 2015. ISSN 2041-1723. doi: 10.1038/ncomms9885. URL https://www.nature.com/articles/ncomms9885.

M. Rigotti, O. Barak, M. R. Warden, X.-J. Wang, N. D. Daw, E. K. Miller, and S. Fusi. The importance of mixed selectivity in complex cognitive tasks. *Nature*, 497(7451):585–590, May 2013. ISSN 1476-4687. doi: 10.1038/nature12160. URL https://www.nature.com/articles/nature12160. Number: 7451.

L. M. Romanski and P. S. Goldman-Rakic. An auditory domain in primate prefrontal cortex. *Nature Neuroscience*, 5(1):15–16, Jan. 2002. ISSN 1097-6256, 1546-1726. doi: 10.1038/nn781. URL https://www.nature.com/articles/nn781.

P. J. Rousseeuw. Silhouettes: A graphical aid to the interpretation and validation of cluster analysis. *Journal of Computational and Applied Mathematics*, 20:53–65, Nov. 1987. ISSN 0377-0427. doi: 10.1016/0377-0427(87)90125-7. URL https://www.sciencedirect.com/science/article/pii/0377042787901257.

S. Ryali, T. Chen, K. Supekar, and V. Menon. A parcellation scheme based on von Mises-Fisher distributions and Markov random fields for segmenting brain regions using resting-state fMRI. *NeuroImage*, 65:83–96, Jan. 2013. ISSN 1053-8119. doi: 10.1016/j.neuroimage.2012.09.067. URL https://www.sciencedirect.com/science/article/pii/S1053811912009858.

A. Schaefer, R. Kong, E. M. Gordon, T. O. Laumann, X.-N. Zuo, A. J. Holmes, S. B. Eickhoff, and B. T. T. Yeo. Local-Global Parcellation of the Human Cerebral Cortex from Intrinsic Functional Connectivity MRI. *Cerebral Cortex*, 28(9):3095–3114, Sept. 2018. ISSN 1047-3211. doi: 10.1093/cercor/bhx179. URL https://doi.org/10.1093/cercor/bhx179.

M. L. Seghier and C. J. Price. Interpreting and Utilising Intersubject Variability in Brain Function. *Trends in Cognitive Sciences*, 22(6):517–530, June 2018. ISSN 1364-6613, 1879-307X. doi: 10.1016/j.tics.2018.03.003. URL https://www.cell.com/trends/cognitive-sciences/abstract/S1364-6613(18)30064-0.

J. M. Shine, P. G. Bissett, P. T. Bell, O. Koyejo, J. H. Balsters, K. J. Gorgolewski, C. A. Moodie, and R. A. Poldrack. The Dynamics of Functional Brain Networks: Integrated Network States during Cognitive Task Performance. *Neuron*, 92(2):544–554, Oct. 2016. ISSN 0896-6273. doi: 10.1016/j.neuron.2016.09.018. URL https://www.sciencedirect.com/science/article/pii/S0896627316305773.

M. Straathof, M. R. Sinke, R. M. Dijkhuizen, and W. M. Otte. A systematic review on the quantitative relationship between structural and functional network connectivity strength in mammalian brains. *Journal of Cerebral Blood Flow & Metabolism*, 39(2):189–209, Feb. 2019. ISSN 0271-678X. doi: 10.1177/0271678X18809547. URL https://www.ncbi.nlm.nih.gov/pmc/articles/PMC6360487/.

T. Sugihara, M. D. Diltz, B. B. Averbeck, and L. M. Romanski. Integration of Auditory and Visual Communication Information in the Primate Ventrolateral Prefrontal Cortex. *Journal of Neuroscience*, 26(43):11138–11147, Oct. 2006. ISSN 0270-6474, 1529-2401. doi: 10.1523/JNEUROSCI.3550-06.2006. URL https://www.jneurosci.org/content/26/43/11138.

B. Thirion, A. Thual, and A. L. Pinho. From deep brain phenotyping to functional atlasing. *Current Opinion in Behavioral Sciences*, 40:201–212, Aug. 2021. ISSN 2352-1546. doi: 10.1016/j.cobeha.2021.05.004. URL https://www.sciencedirect.com/science/article/pii/S2352154621001121.

B. T. Thomas Yeo, F. M. Krienen, J. Sepulcre, M. R. Sabuncu, D. Lashkari, M. Hollinshead, J. L. Roffman, J. W. Smoller, L. Zöllei, J. R. Polimeni, B. Fischl, H. Liu, and R. L. Buckner. The organization of the human cerebral cortex estimated by intrinsic functional connectivity. *Journal of Neurophysiology*, 106(3):1125–1165, Sept. 2011. ISSN 0022-3077, 1522-1598. doi: 10.1152/jn.00338.2011. URL https://www.physiology.org/doi/10.1152/jn.00338.2011.

D. Y. Tsao, N. Schweers, S. Moeller, and W. A. Freiwald. Patches of face-selective cortex in the macaque frontal lobe. *Nature Neuroscience*, 11(8):877–879, Aug. 2008. ISSN 1546-1726. doi: 10.1038/nn.2158. URL https://www.nature.com/articles/nn.2158.

J. D. Xiang, J. C. Martinez-Trujillo, T. W. Schmitz, and M. Mur. Toward task mapping of primate prefrontal cortex. *Neuropsychologia*, 219:109234, Dec. 2025a. ISSN 0028-3932. doi: 10.1016/j.neuropsychologia.2025.109234. URL https://www.sciencedirect.com/science/article/pii/S0028393225001691.

J. D. Xiang, M. Roussy, B. Corrigan, R. A. Gulli, R. Luna, M. H. Mofrad, L. Muller, J. Diedrichsen, T. W. Schmitz, J. Martinez-Trujillo, and M. Mur. Task-specific topographic maps of neural activity in the primate lateral prefrontal cortex, July 2025b. URL https://www.biorxiv.org/content/10.1101/2024.05.10.591729v2. ISSN: 2692-8205 Pages: 2024.05.10.591729 Section: New Results.

R. Xu, N. P. Bichot, A. Takahashi, and R. Desimone. The cortical connectome of primate lateral prefrontal cortex. *Neuron*, 110(2):312–327.e7, Jan. 2022. ISSN 0896-6273. doi: 10.1016/j.neuron.2021.10.018. URL https://www.sciencedirect.com/science/article/pii/S0896627321007868.

T. Xu, A. Opitz, R. C. Craddock, M. J. Wright, X.-N. Zuo, and M. P. Milham. Assessing Variations in Areal Organization for the Intrinsic Brain: From Fingerprints to Reliability. *Cerebral Cortex*, 26(11):4192–4211, Oct. 2016. ISSN 1047-3211. doi: 10.1093/cercor/bhw241. URL https://doi.org/10.1093/cercor/bhw241.

E. Yacoub, N. Harel, and K. Uğurbil. High-field fMRI unveils orientation columns in humans. *Proceedings of the National Academy of Sciences*, 105(30):10607–10612, July 2008. doi: 10.1073/pnas.0804110105. URL https://www.pnas.org/doi/10.1073/pnas.0804110105.

G. R. Yang, M. R. Joglekar, H. F. Song, W. T. Newsome, and X.-J. Wang. Task representations in neural networks trained to perform many cognitive tasks. *Nature Neuroscience*, 22(2): 297–306, Feb. 2019. ISSN 1097-6256, 1546-1726. doi: 10.1038/s41593-018-0310-2. URL http://www.nature.com/articles/s41593-018-0310-2.

D. Zhi, M. King, C. R. Hernandez-Castillo, and J. Diedrichsen. Evaluating brain parcellations using the distance-controlled boundary coefficient. *Human Brain Mapping*, 43(12):3706–3720, 2022. ISSN 1097-0193. doi: 10.1002/hbm.25878. URL https://onlinelibrary.wiley.com/doi/abs/10.1002/hbm.25878. _eprint: https://onlinelibrary.wiley.com/doi/pdf/10.1002/hbm.25878.

D. Zhi, L. Shahshahani, C. Nettekoven, A. L. Pinho, D. Bzdok, and J. Diedrichsen. A hierarchical Bayesian brain parcellation framework for fusion of functional imaging datasets. *Imaging Neuroscience*, 3:imag_a_00408, Jan. 2025. ISSN 2837-6056. doi: 10.1162/imag_a_00408. URL https://doi.org/10.1162/imag_a_00408.
